# Regulated assembly of the ER membrane protein complex

**DOI:** 10.1101/2020.07.20.213066

**Authors:** Tino Pleiner, Kurt Januszyk, Giovani Pinton Tomaleri, Robert S. Oania, Masami Hazu, Michael J. Sweredoski, Annie Moradian, Alina Guna, Rebecca M. Voorhees

## Abstract

The assembly of nascent proteins into multi-subunit complexes is tightly regulated to maintain cellular homeostasis. The ER membrane protein complex (EMC) is an essential insertase that requires seven membrane-spanning and two soluble subunits for function. Here we show that the kinase With no lysine 1 (WNK1), known for its role in hypertension and neuropathy, is required for assembly of the human EMC. WNK1 uses a conserved amphipathic helix to stabilize the soluble subunit, EMC2, by binding to the EMC2-8 interface. Shielding this hydrophobic surface prevents promiscuous interactions of unassembled EMC2 and precludes binding of ubiquitin ligases, permitting assembly. Using biochemical reconstitution, we show that after EMC2 reaches the membrane, its interaction partners within the EMC displace WNK1, and similarly shield its exposed hydrophobic surfaces. This work describes an unexpected role for WNK1 in protein biogenesis, and defines the general requirements of an assembly factor that will apply across the proteome.

## Introduction

Protein complex assembly is a major challenge within the crowded cellular environment. Nearly half of the eukaryotic proteome is organized into multi-subunit complexes, and when unassembled these subunits are prone to non-productive interactions and aggregation (Marsh and Teichmann, 2015; Juszkiewicz and Hegde, 2018). To prevent cytotoxicity, complexes either assemble co-translationally (Duncan and Mata, 2011; Shiber et al., 2018), or rely on specific chaperones to coordinate their assembly (Jäkel et al., 2002; Kihm et al., 2002). Given the enormous diversity of protein subunits, the suite of factors required to regulate protein complex assembly is incompletely defined.

One type of multiprotein complex that faces additional challenges for assembly are those that contain both soluble and membrane-spanning subunits, which require the temporal and spatial coordination of protein synthesis and membrane insertion between the cytosol and endoplasmic reticulum (ER). These include complexes such as the calcium and potassium voltage-gated ion channels, as well as the ubiquitously expressed ER membrane protein complex (EMC) (Jonikas et al., 2009; Wideman, 2015).

The EMC is an evolutionarily conserved ER-resident complex, which plays a critical role in the biogenesis of a diverse set of membrane proteins (Chitwood and Hegde, 2019). In mammals, the EMC is composed of seven integral membrane subunits (EMC1, 3-7, and 10) and two soluble cytosolic subunits (EMC2 and 8), the majority of which are required for both its stability and function (Figures 1A and S1A). Detailed biochemical reconstitution experiments have shown that the EMC can both co-translationally insert TMDs in the N_exo_-topology, and post-translationally insert tail-anchored proteins with TMDs of moderate hydrophobicity (Chitwood et al., 2018; Guna et al., 2018). Proteomic interrogation of EMC function also suggests it plays a role in stabilizing multi-pass membrane proteins, including many viral coat proteins (Barrows et al., 2019; Chitwood and Hegde, 2019; Ngo et al., 2019; Shurtleff et al., 2018; Tian et al., 2019).

**Figure 1.**
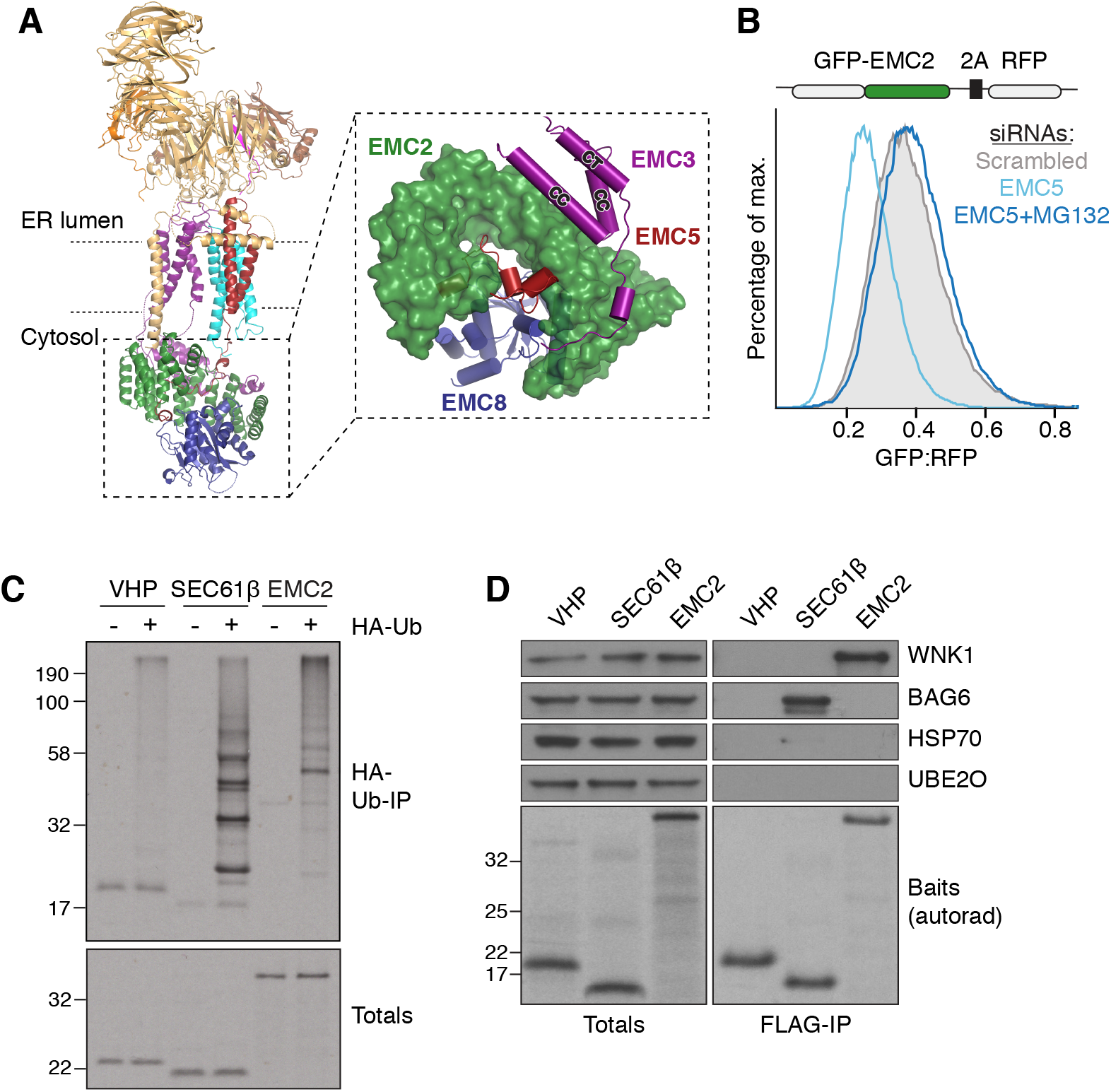
Unassembled EMC2 binds WNK1. (**A**) Overview of the cytosolic domain of the human EMC, assembled around EMC2 (PDB 6WW7; Pleiner et al., 2020). Inset: close-up on the cytosolic domain as viewed from the membrane. (**B**) HEK293T cells stably expressing GFP-EMC2 were treated with either scrambled (gray) or EMC5 siRNA (blue) −/+ the proteasome inhibitor MG132. The relative levels of GFP-EMC2 normalized to an internal RFP expression control (described in methods) are plotted as a histogram and reflect changes in post-translational subunit stability. (**C**) EMC2 is ubiquitinated in vitro. ^35^S-methionine-labeled villin headpiece (VHP; an inert control), the membrane protein SEC61β (previously shown to be robustly ubiquitinated when expressed in the absence of membranes; Hessa et al., 2011), and EMC2 were translated in rabbit reticulocyte lysate (RRL) in the presence of HA-tagged ubiquitin. Ubiquitinated proteins were immunoprecipitated via the HA-tag and analyzed by SDS-PAGE and autoradiography. (**D**) Native immunoprecipitation of ^35^S-methionine labeled 3xFLAG-tagged VHP, SEC61β and EMC2 from rabbit reticulocyte lysate (RRL). Eluates were analyzed by SDS-PAGE and Western blotting with the indicated antibodies. Baits were detected by autoradiography (autorad).

The EMC is therefore an essential cellular machine and previous work suggests that its assembly is highly regulated (Volkmar et al., 2019). The cell must ensure that the nine subunits of the EMC are assembled at the correct stoichiometry, while preventing inappropriate interactions of isolated subunits or assembly intermediates during this process. Given the high abundance of the EMC and its importance in human cells (Kulak et al., 2014), we postulated that dedicated regulatory machinery may be involved. Because the soluble subunits, such as EMC2, may be particularly reliant on assembly factors for transport to the ER, we interrogated the mechanism of EMC assembly focusing on its cytosolic domain.

## Results

### Orphan EMC2 is unstable and binds WNK1

Previous work established that the stability of the EMC subunits is interdependent; orphan subunits that cannot assemble or are synthesized in excess are degraded post-translationally by the ubiquitin-proteasome pathway (Figure S1B) (Volkmar et al., 2019). This interdependence is especially evident for EMC2, a soluble component of the complex that is robustly targeted for degradation both in cells and *in vitro* when unable to assemble (Figures 1B-C and S1C). The structure of the human EMC provides an explanation for the short half-life of orphan EMC2: EMC2 is a superhelical architectural scaffold, responsible for anchoring both the soluble and membrane spanning subunits of the complex (Figure 1A) (Pleiner et al., 2020). Thus, unassembled EMC2 has multiple exposed hydrophobic interfaces that ultimately interact with other EMC subunits.

We rationalized that EMC2 could require a novel assembly factor because the levels of free EMC2 are tightly regulated in cells, leading to little or no detectable unassembled EMC2. Further, exogenously expressed EMC2 is soluble and monodisperse (Figure S1D), and did not recruit nonspecific chaperones such as Hsp70, or known assembly or quality control factors such as UBE2O and BAG6 (Figure 1D) (Hessa et al., 2011; Yanagitani et al., 2017).

To identify putative cytosolic binding partners, *in vitro* translated EMC2 was affinity-purified under native conditions from rabbit reticulocyte lysate (RRL) and analyzed by mass spectrometry. Using this strategy, we identified the kinase With no Lysine 1 (WNK1) (Xu et al., 2000) as a specific interaction partner of EMC2 both *in vitro* and in cells, and validated the interaction by immunoblotting (Figures 1D and S1E). Given the relatively low expression level of EMC2 in RRL (~30 nM), we concluded that WNK1 is a selective and relatively stable interaction partner of EMC2.

WNK1 is a ubiquitously expressed regulator of cellular ion homeostasis and cell volume (Alessi et al., 2014; Hadchouel et al., 2016; Shekarabi et al., 2017). Mutations in the WNK1 gene cause disordered renal ion transport leading to hereditary hypertension (Wilson et al., 2001), as well as defects in sensory and autonomic neurons that lead to peripheral neuropathy (Shekarabi et al., 2008). During osmotic stress, WNK1 initiates a kinase cascade to activate ion channels such as the Na-K-Cl cotransporter (NKCC1) to re-establish cellular ion levels. Though WNK1 has never previously been implicated in protein assembly, its pleiotropic phenotypes, particularly as related to a diversity of membrane proteins, could be consistent with a general role in membrane protein biogenesis. In fact, many previously described EMC substrates regulate ion homeostasis and cell volume (Table S1) (Shurtleff et al., 2018; Tian et al., 2019). Interestingly, we found that one of WNK1’s primary downstream targets, NKCC1, also relies on EMC for biogenesis (Figures S1F and S1G). We thus investigated a potential role for WNK1 in EMC assembly.

### WNK1 is an EMC assembly factor

We reasoned that a bona fide EMC2 assembly factor would: (i) transiently stabilize unassembled EMC2; (ii) be required for the assembly and function of the entire EMC; and (iii) directly interact with unassembled, but not assembled EMC2. First, we used a quantitative flow cytometry assay to measure the stability of GFP-EMC2, using RFP as an internal expression control. We found that three independent small interfering RNAs (siRNAs) targeting WNK1 destabilized GFP-EMC2 (Figures 2A and S2A). The effect could be rescued by expression of an siRNA resistant WNK1, excluding off-target effects (Figure 2A). Thus, we concluded that WNK1 stabilizes unassembled EMC2 and meets the first criterion for an assembly factor.

**Figure 2.**
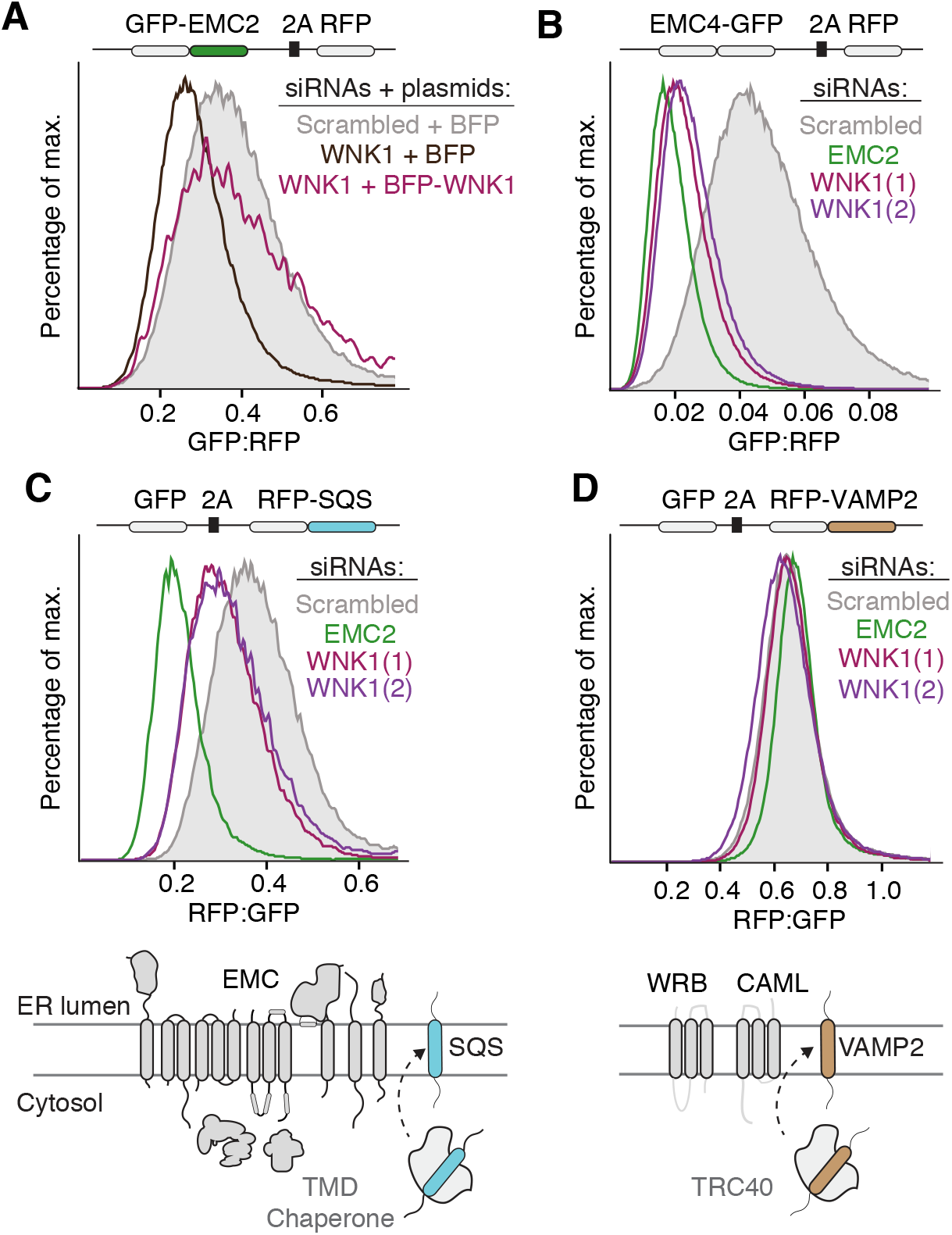
WNK1 is an EMC assembly factor. (**A**) HEK293T cells stably expressing GFP-EMC2 were transiently transfected with plasmids encoding either BFP or BFP-tagged siRNA-resistant WNK1. Cells were treated with either scrambled or WNK1 siRNA, and BFP-positive cells were analyzed by flow cytometry. The relative levels of GFP-EMC2, normalized to an internal RFP expression control (described in methods), are plotted as a histogram and reflect changes in subunit stability. (**B-D**) Scrambled or WNK1 siRNA treatment of EMC4-GFP, RFP-squalene synthase (SQS; a tail-anchored EMC substrate) and RFP-VAMP2 (a tail-anchored WRB-CAML substrate) cell lines and analysis by flow cytometry.

Using a similar assay, we expressed EMC4-GFP and EMC5-GFP, and verified their incorporation into the intact complex (Figure S2B). WNK1 depletion also destabilized these membrane-embedded EMC subunits, phenocopying EMC2 knockdown (Figures 2B and S2C). Importantly, the observed decrease in EMC levels upon WNK1 knockdown also compromised EMC function. We demonstrated that treatment with several independent WNK1 siRNAs caused biogenesis defects for the post- and co-translational EMC substrates, SQS and OPRK1, but had no effect on the EMC independent substrates, VAMP2 and TRAM2 (Figures 2C-D and S2D). These results exclude the possibility of a general dysregulation of the ER and suggest a specific role for WNK1 in the assembly of the EMC. Additionally, the pan-WNK kinase inhibitor WNK463 (Yamada et al., 2016) had no effect on EMC subunit or substrate levels, suggesting that WNK1’s role in assembly does not require its kinase function (Figure S3). WNK1 is thus required for assembly of the intact EMC and meets the second criterion for an assembly factor.

Binding assays with purified recombinant components confirmed that EMC2 and WNK1 interact directly and form a stable complex *in vitro* (Figure 3A). Truncation analysis identified a single twenty-six amino acid region of WNK1 (amino acids 2241-2266) that was both necessary and sufficient for binding to EMC2 (Figure 3A). This previously unannotated sequence in the C-terminus of WNK1 is predicted to form a conserved amphipathic helix in metazoans (Figure S4A) and is sufficient to rescue the destabilizing effect of WNK1 knockdown on EMC2 levels in cells (Figure 3B). This amphipathic sequence is conserved amongst paralogs of WNK1, and the analogous regions in WNK2 and 3 also bind to EMC2 *in vitro* (Figures S4A-B). Finally, experiments comparing binding of WNK1-helix to either free EMC2 or purified EMC (normalized to EMC2 levels), confirmed that the WNK1-helix selectively interacts with unassembled EMC2 (Figure 3C). We thus determined that WNK1 meets all the criteria of an EMC2 assembly factor.

**Figure 3.**
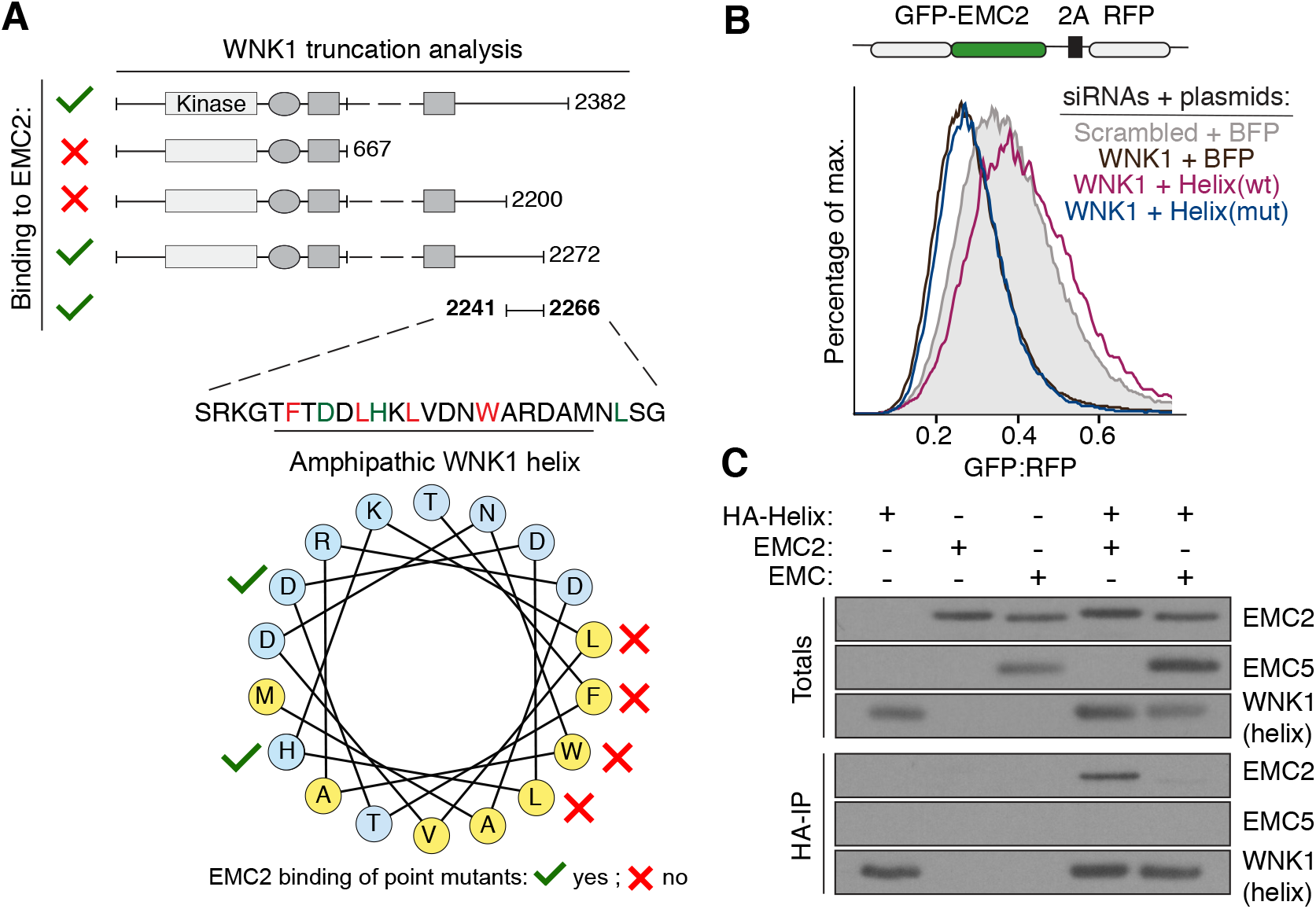
A conserved amphipathic α-helix in WNK1 binds to unassembled EMC2. (**A**) FLAG-tagged WNK1 and its truncations were tested for binding to EMC2 (see methods). The EMC2 binding site was mapped to a previously uncharacterized C-terminal peptide that is predicted to form a conserved amphipathic α-helix (see Figure S4). Polar residues are shown in blue and hydrophobic residues in yellow. (**B**) HEK293T cells stably expressing GFP-EMC2 were transiently transfected with plasmids encoding either BFP or BFP-tagged wild type (wt) or L2250K mutant (mut) WNK1 helix. Cells were treated with either scrambled or WNK1 siRNA and analyzed by flow cytometry as in Figure 2A. (**C**) Intact EMC, purified from stable EMC5-3xFLAG HEK293T cells, and recombinant EMC2 were incubated with 3xHA-tagged WNK1 helix and immunoprecipitated with anti-HA resin. Totals and eluates were analyzed by Western blotting with the indicated antibodies.

### The amphipathic WNK1 helix shields the hydrophobic EMC2-8 interface

To understand why EMC2, a soluble and apparently monodisperse protein, requires an assembly factor, we defined the molecular nature of the EMC2-WNK1 interaction. Mutational analysis of the WNK1-helix suggested that binding to EMC2 is mediated via the hydrophobic face of the helix (Figure S4C). Point mutations to F2246, L2250, L2253, or W2257, but not D2248 or H2251, disrupted binding *in vitro* and abolished the stabilizing effect of the wild type helix on exogenously overexpressed EMC2 in cells (Figure 4A). Site-specific crosslinking using a protein with an exposed transmembrane domain confirmed that unassembled EMC2 is able to form nonspecific interactions with hydrophobic peptides (Figure 4B). Binding of wild type, but not mutant WNK1-helix, competed for crosslinking with this substrate, consistent with a role for WNK1 in shielding this exposed surface of EMC2 prior to assembly. Therefore, one function of WNK1 is to prevent promiscuous interactions of an unprotected hydrophobic patch on unassembled EMC2.

**Figure 4.**
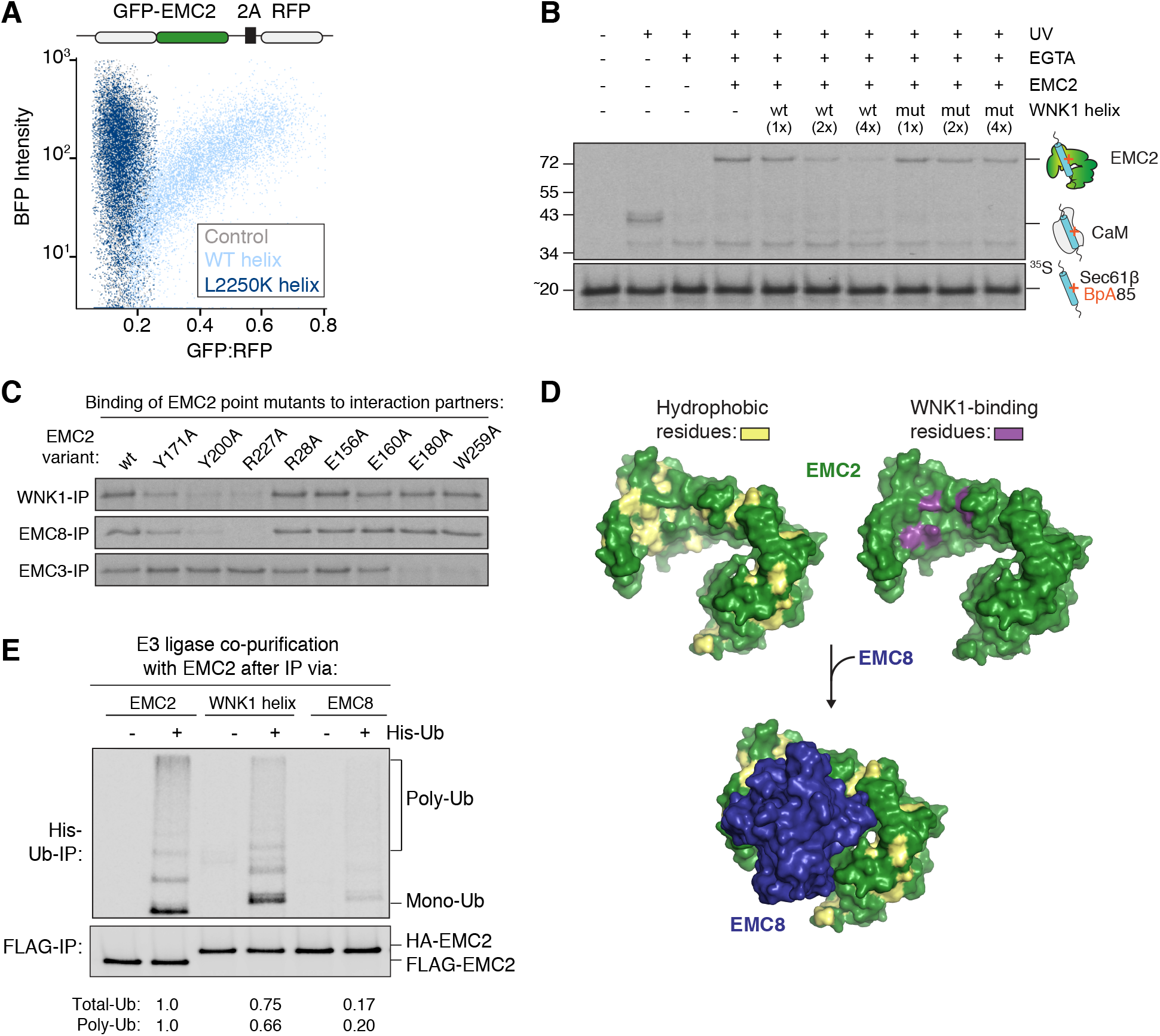
WNK1 binds the EMC2-EMC8 interface and stabilizes unassembled EMC2. (**A**) HEK293T cells stably expressing GFP-EMC2 were transfected with plasmids encoding BFP, BFP-tagged WNK1 wild type (wt) or L2250K mutant helix. Cells were analyzed by flow cytometry and their relative GFP intensity, normalized to an internal RFP expression control, was plotted against BFP intensity. (**B**) An ^35^S-methionine-labeled representative hydrophobic protein (SEC61β) was produced with the BpA photo–crosslinker within the TMD as a complex with calmodulin (CaM) in the PURE *in vitro* translation system. The CaM-SEC61β complexes were incubated with EMC2 alone or pre-formed complexes of EMC2 with increasing concentrations of purified wild type (wt) or L2250K mutant WNK1 helix. Except for the -UV control, all reactions were irradiated and analyzed by SDS-PAGE and autoradiography. (**C**) ^35^S-methionine labeled wild type EMC2, or the indicated point mutants were translated in RRL and tested for binding to 3xFLAG-tagged WNK1, EMC8 or EMC3 via co-immunoprecipitation using anti-FLAG resin. EMC2 mutants that are unable to bind WNK1 also do not bind EMC8, but are capable of interacting with EMC3, indicating proper folding. (**D**) Depicted is the surface representation of EMC2 (PDB 6WW7; Pleiner et al., 2020) in which hydrophobic residues have been highlighted in yellow (left) or mutations that affected WNK1 binding in purple (right). The WNK1 binding site overlapped with the binding site of EMC8 (blue), which is characterized by an extended hydrophobic patch. EMC2 is viewed from the cytosol. (**E**) ^35^S-methionine labeled EMC2 was translated in RRL and purified either directly via a 3xFLAG-tag or indirectly via 3xFLAG-tagged WNK1 helix or EMC8 (translation of 3xHA-tagged EMC2). The purified complexes were analyzed for ubiquitination competence to detect the presence of co-purifying E3 ubiquitin ligases. Following enrichment for ubiquitinated species, the total intensity of ubiquitinated species in all samples was quantified and normalized to the 3xFLAG-EMC2 sample.

In considering the potential location of WNK1’s binding site, we were struck by the presence of a conserved hydrophobic interface between EMC2 and EMC8 (Pleiner et al., 2020). Using purified recombinant components, we found that the interaction of EMC2 with WNK1-helix is mutually exclusive to the interaction with EMC8 (Figure S5). Structure-guided mutagenesis confirmed that the WNK1 recognition site partially overlaps with that of EMC8, as mutations that disrupted WNK1 binding also decreased EMC8 binding (Figures 4C-D). WNK1 thus directly shields the interface of EMC2 that is eventually occupied by EMC8, analogous to the role of AHSP in the assembly of adult hemoglobin (Kihm et al., 2002). Comparison of the human and *S. cerevisiae* EMC structures suggests that in yeast, which lack an EMC8 homolog, the analogous WNK1 binding site is occupied by the N-terminus of EMC4 upon assembly (Figure S5D) (Bai et al., 2020).

We reasoned that given its hydrophobic nature, the EMC8 and WNK1 binding site could also serve as a recognition site for quality control factors when exposed. To test this, we immunoprecipitated EMC2 under native conditions from RRL either on its own, or when bound to WNK1 or EMC8 (Figure 4E). When purified on its own, EMC2 became robustly modified with His-tagged ubiquitin after incubation with recombinant E1 and E2 enzymes, indicating co-purification of an endogenous E3 ubiquitin ligase. In contrast, binding to WNK1 or EMC8 partially or almost completely diminished the subsequent ubiquitination of EMC2, respectively. WNK1 therefore stabilizes EMC2 during assembly by competing with E3 ubiquitin ligase binding to the EMC2-8 subunit interface.

### Reconstitution of the assembly pathway for the EMC cytosolic domain

In order to position WNK1 in the EMC assembly pathway, we determined the hierarchy of EMC subunit interactions. In line with its central role in the stability of the EMC, EMC2 serves as a scaffold for anchoring both the soluble and membrane-spanning subunits of the complex, including EMC1, 3, 5, 4 and 8 (Bai et al., 2020; Pleiner et al., 2020). *In vitro* translation of EMC2 and 3 in the presence of canine rough microsomes established that EMC3 is sufficient to recruit EMC2 to the membrane (Figure 5A). Experiments using recombinant components revealed that EMC3 binding requires both the coiled-coil and complete C-terminus of EMC3, and is compatible with formation of a trimeric complex with either WNK1 or EMC8 (Figures 5B and S6A).

**Figure 5.**
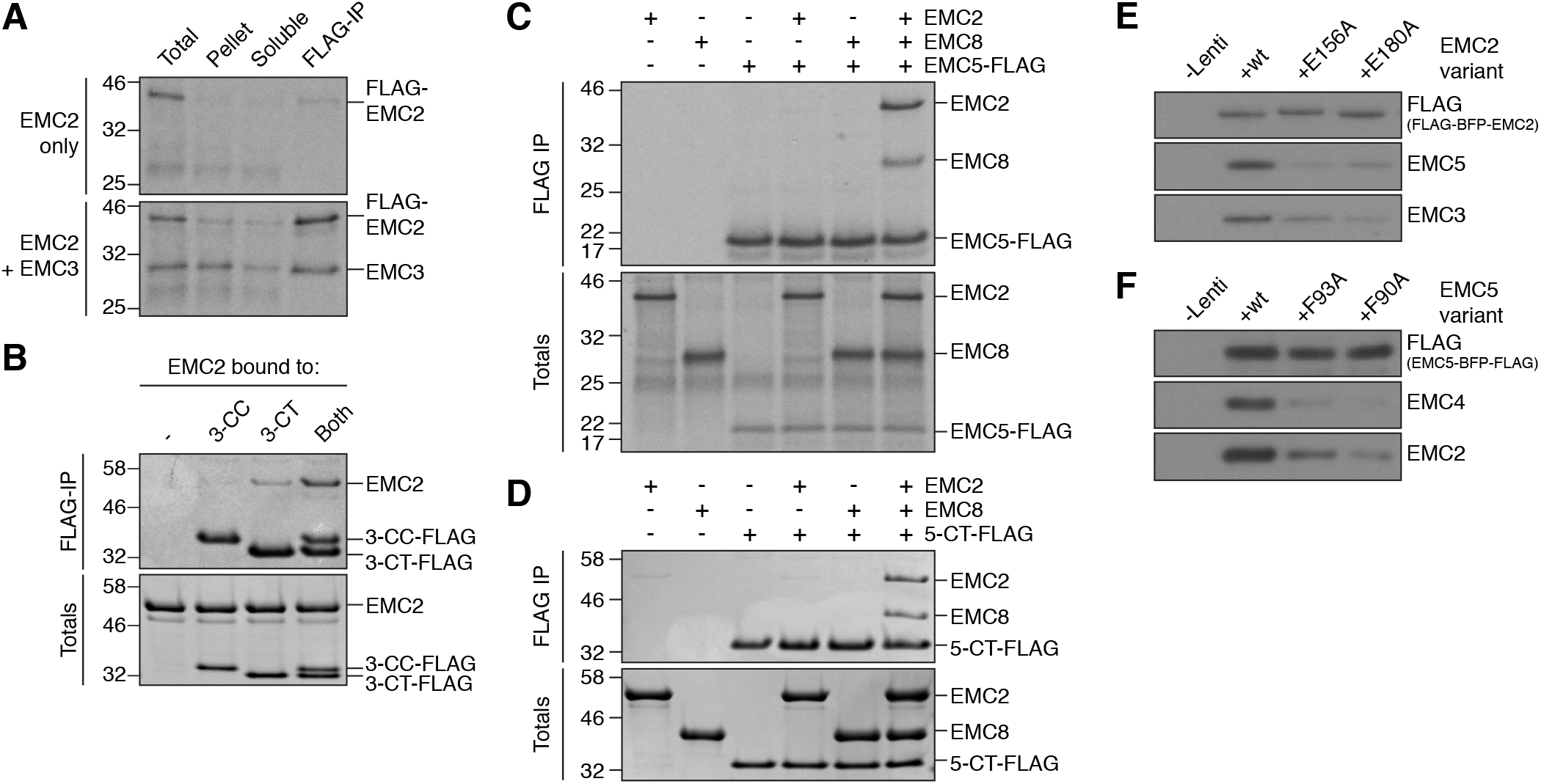
*In vitro* assembly of the cytosolic domain of the EMC. (**A**) ^35^S-methionine-labeled 3xFLAG-EMC2 was translated alone or with untagged EMC3 in RRL in the presence of canine rough microsomes (cRMs). cRMs were isolated by centrifugation through a sucrose cushion, resuspended, and solubilized in digitonin. The resulting detergent lysate was subjected to FLAG-IP and analyzed by SDS-PAGE and autoradiography. (**B**) Both the EMC3 coiled coil (3-CC) and C-terminus (3-CT) are needed to interact with EMC2. Purified 3xFLAG-tagged 3-CC and 3-CT were incubated with EMC2 either alone or in combination and then subjected to FLAG-IP. Eluates were analyzed by SDS-PAGE and Coomassie staining. (**C**) Binding assays were performed as in (A) for full-length 3xFLAG-tagged EMC5 alone or in combination with 3xHA-tagged EMC2, EMC8, or both. (**D**) EMC5 C-terminus (5-CT) is sufficient to bind the EMC2-8 complex. 3xFLAG-tagged 5-CT was incubated either with buffer, EMC2, EMC8, or both and subjected to native FLAG-IP. Totals and eluates were analyzed by SDS-PAGE and Coomassie staining. (**E**) HEK293T cells stably expressing exogenous 3xFLAG-tagged wild type or mutant EMC2 were lysed and subjected to native FLAG-IP. Eluates were analyzed by Western blotting to determine the efficiency of assembly of wild type and mutant EMC2. (**F**) As in E, but with wild type or mutant 3xFLAG-tagged EMC5.

In contrast, EMC2 could only bind EMC5 in the presence of EMC8, but not WNK1, suggesting that EMC8 binding must precede interaction with EMC5 (Figures 5C-D and S6B). In yeast, which lacks an EMC8 homolog, EMC2 binds EMC5 independently (Figures S6C-D). As EMC8 forms no direct contacts to the membrane-spanning EMC subunits, we postulate that EMC8 displaces WNK1 in the cytosol. WNK1 binding therefore precedes EMC8 binding to EMC2. This is supported by a preferential accumulation of EMC2 in complex with WNK1, rather than EMC8, upon proteasome inhibition (Figures S6E-F).

After WNK1 displacement, the heterodimeric EMC2-8 complex is recruited to the membrane by interaction with the EMC3 and 5 cytosolic loops. Mutations to EMC2 at the EMC2-5 and EMC2-3 interface decreased incorporation of EMC2 into the complex both *in vitro* and in cells, suggesting that membrane tethering by both EMC5 and EMC3 is critical for assembly (Figures 5E, S6G-I). Similarly, mutations to the C-terminus of EMC5 that prevent EMC2 interaction also markedly decreased EMC5 assembly into the intact EMC (Figures 5F and S6G). Using this information, we reconstituted the cytosolic domain of the EMC, including EMC2 and 8 and the loops of EMC3 and 5, with purified, recombinant components and verified it forms a stable tetrameric complex that does not interact with WNK1 (Figure 6A).

**Figure 6.**
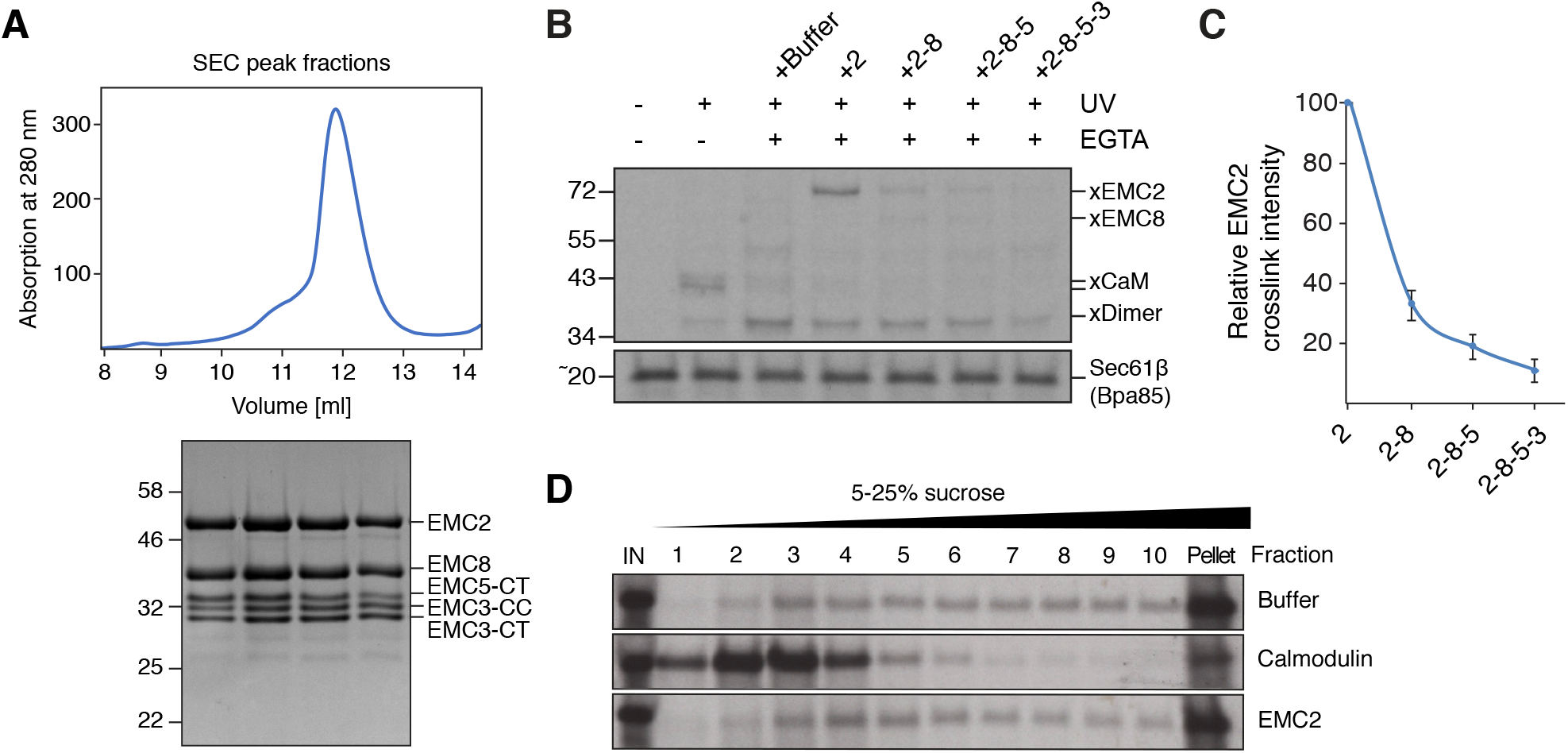
The reconstituted EMC cytosolic domain does not stably interact with substrates. (**A**) Reconstitution of the EMC cytosolic domain from purified components: EMC2, 8, 3-CC, 3-CT and 5-CT. The complex was analyzed by size-exclusion chromatography (SEC), followed by SDS-PAGE and Coomassie staining of the peak fractions. (**B**) A complex of calmodulin (CaM) and ^35^S-methionine-labeled SEC61β (a tail-anchored EMC substrate), containing the photocrosslinker BpA in the TMD, was generated in the PURE *in vitro* translation system. The CaM-SEC61β complexes were incubated with either purified EMC2 alone or complexes of EMC2 with the indicated binding partners. After UV irradiation the samples were analyzed by SDS-PAGE and autoradiography. (**C**) Quantification of the EMC2-SEC61β crosslink intensity from (B) (n=3). (**D**) ^35^S-methionine-labeled SQS (a tail-anchored EMC substrate) was translated in the PURE system in the presence or absence of 12 *μ*M calmodulin or 12 *μ*M EMC2. Translation reactions were separated by native size through a sucrose gradient, and fractions were analyzed by SDS-PAGE and autoradiography. Fraction 11 represents the resuspended pellet.

Reconstitution of the intact cytosolic domain permitted interrogation of its potential role in substrate interaction. It was recently hypothesized that EMC2 contains a substrate-binding groove that could potentially accommodate an unfolded TMD during its transit into the membrane (O’Donnell et al., 2020). This model was based on experiments performed using the EMC2-8 sub-complex; however analysis of the complete EMC structure indicates that the proposed groove is occluded by the C-terminus of EMC5, which would have to be displaced to allow substrate engagement (Pleiner et al., 2020). Using the intact, tetrameric cytosolic domain, we found that EMC8 binding strongly reduced crosslinking of TMDs to unassembled EMC2 (Figures 6B-C). Addition of the EMC5 C-terminus, and to a lesser extent the cytosolic loops of EMC3, further reduced crosslinking intensity. Thus analogous to WNK1, EMC2’s binding partners collectively reduce non-specific interactions by shielding the hydrophobic interfaces of EMC2 upon assembly. Furthermore, in contrast to calmodulin, a TMD chaperone with an ordered hydrophobic groove, EMC2 was not able to solubilize an *in vitro* translated transmembrane domain (Figure 6D). Taken together, these data suggest EMC2 does not stably bind substrates.

## Discussion

We have demonstrated that in addition to its established role in regulating ion homeostasis, WNK1 is an assembly factor for the EMC (Figure 7). WNK1 uses a conserved amphipathic helix to shield an exposed subunit interface on EMC2, preventing nonspecific interactions and recognition by quality control factors to allow assembly. Mutational analysis of the WNK1 helix further suggested that it is required for the previously observed recruitment of the kinase to clathrin-coated vesicles during osmotic stress (Figure S7) (Zagórska et al., 2007). This unexpected finding suggests an intriguing connection between protein biogenesis in the ER and osmotic stress.

**Figure 7.**
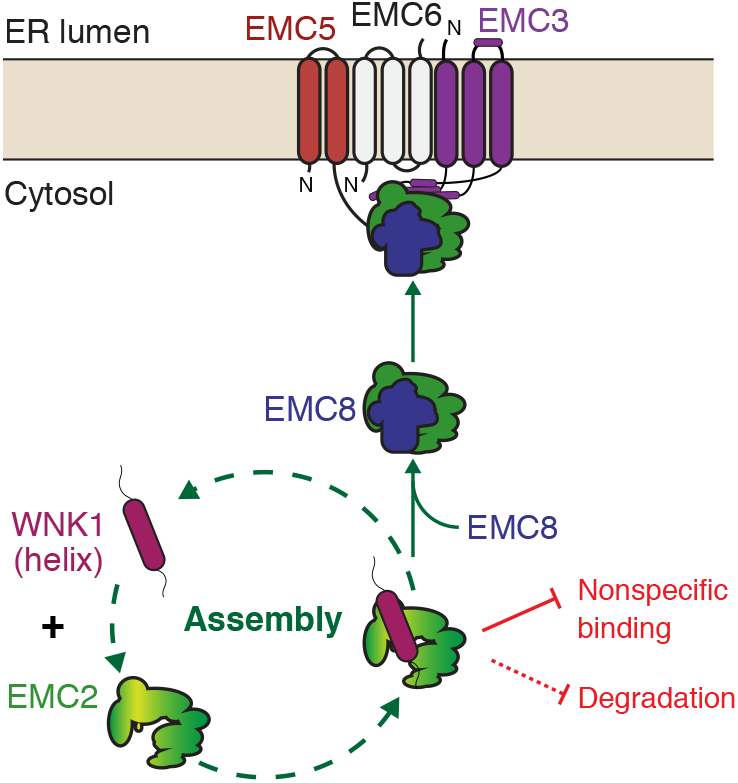
Model for assembly of the cytosolic domain of the EMC. WNK1 stabilizes unassembled EMC2 by using a conserved C-terminal amphipathic α-helix to prevent nonspecific binding and premature degradation in the cytosol. EMC8 then displaces WNK1 in the cytosol, allowing assembly with the membrane-bound subunits EMC3 and EMC5.

By investigating EMC2 assembly, we can now analyze the functional role of the EMC cytosolic domain, and extract generalizable principles for multi-subunit protein complex assembly. By analogy to the relationship between TRC40 and WRB, it was speculated that EMC2, which is anchored at the membrane by the coiled-coil of EMC3 (homologous to WRB), could be involved in capturing substrates in the cytosol. However unlike TRC40, which has an ordered methionine-rich TMD binding groove (Anghel et al., 2017; Mariappan et al., 2011; Mateja et al., 2015), recent structural analysis reveals that when assembled, EMC2 does not contain an obvious hydrophobic groove in either yeast or humans (Bai et al., 2020; Pleiner et al., 2020).

Recently, experiments utilizing the EMC2-8 heterodimer identified a hydrophobic cleft that was proposed to accommodate an unfolded TMD (O’Donnell et al., 2020). Crosslinking experiments with the 2-8 heterodimer suggested that this sub-complex could transiently interact with substrate, and mutations to the proposed hydrophobic cleft at the EMC2-5 interface affected EMC insertion. However, we found that in the context of the reconstituted cytosolic domain (including EMC3, 5, and 8) EMC2 does not appreciably crosslink to TMDs (Figures 6B-C), and thus at least *in vitro*, a substrate cannot efficiently displace the C-terminus of EMC5. Further, mutations to the EMC2-5 interface prevented assembly in cells (Figures 6E-F and S6I), which could provide an alternative explanation for the observed insertion defect. Thus, while we do not exclude the possibility that large-scale rearrangements could expose a TMD-binding site in EMC2, our data suggest EMC2 functions primarily as an architectural scaffold rather than a substrate interaction site. In contrast to the substrates of the TRC40-WRB/CAML pathway, the low-hydrophobicity substrates of the EMC are less prone to aggregation and therefore may not require an ordered cytosolic binding groove within the EMC for insertion.

Aside from function, mechanistic details of EMC assembly also shed a new light on WNK1 function. Based on the requirement of WNK1 for assembly of the EMC, many of the pleiotropic phenotypes attributed to WNK1 knockdown or knockout may have to be re-evaluated. In particular, the reported role of WNK1 in surface expression of many membrane proteins could be at least partially explained by the indirect effect of WNK1 depletion on the biogenesis of any EMC-dependent substrate.

We further find that the WNK1 paralogues WNK2 and WNK3 similarly bind to EMC2 and could thus be functionally redundant to WNK1 in mediating EMC assembly. Their expression level, however, and hence potential functional contribution, varies widely among different cell types: WNK1 appears to be the most ubiquitous and highest expressed member of the WNK family (Veríssimo and Jordan, 2001). Lower eukaryotes, which do not express a homolog of EMC8, also do not express a homolog of any WNK family member. However, given that in yeast, the N-terminus of EMC4 occupies the analogous WNK1 binding site on EMC2 upon assembly, it is possible that the requirement for an EMC2 assembly factor is not metazoan-specific.

The EMC2-WNK1 interaction is notably weaker and likely to bury less surface area than the EMC2-8 interaction (1393 Å^2^) (Pleiner et al., 2020), explaining why WNK1 can be displaced upon assembly. A modest affinity of the EMC2-WNK1 complex might be functionally important in order to allow the ultimate triage of terminally unassembled EMC2 for degradation, as has been observed for other biogenesis pathways (Hessa et al., 2011; Yanagitani et al., 2017). These general principles are likely conserved features of assembly factors.

Apart from the EMC, we envision that other protein complexes with subunits that are synthesized and assembled at distinct locations might require novel assembly factors. For example, a major class of membrane protein complexes with soluble subunits are the voltage-gated calcium and potassium ion channels (Buraei and Yang, 2010; Pongs and Schwarz, 2010). Indeed, previous data suggests that like EMC2, some of these soluble subunits are also degraded in the absence of their membrane-embedded binding partners (Menegola and Trimmer, 2006). Further, given the complexity of the membrane-spanning subunits of the EMC, and increasing evidence that membrane protein complex assembly is highly regulated (Inglis et al., 2020), it is likely that the EMC may also rely on dedicated intramembrane chaperones for assembly.

Thus, given the enormous diversity of subunits whose assembly must be tightly regulated by the cell, the known set of assembly factors is undoubtedly incomplete. The strategies reported here lay out a roadmap for their identification and validation. Characterization of these assembly factors will reveal how the cell ensures the stoichiometric assembly of intricate molecular machines in the crowded cellular milieu, and provide insight into how these processes may fail leading to disease (Harper and Bennett, 2016; Oromendia and Amon, 2014).

**Table 1.** EMC substrates involved in ion homeostasis/ cell volume regulation^a^.

| **Gene** | **Protein** | **Reference** |
| --- | --- | --- |
| SLC4A2 | Anion Exchanger 2 (AE2) | (Shurtleff et al., 2018; Tian et al., 2019) |
| SLC6A6 | Taurine transporter (TauT) |  |
| ANO6 | Anoctamine 6/ Small-conductance calcium-activated nonselective cation channel |  |
| CLCN3 | H(+)/Cl(-) exchange transporter 3 | (Tian et al., 2019) |
| LRRC8E | Volume-regulated anion channel subunit LRRC8E |  |
| SLC9A7 | Sodium/hydrogen exchanger 7 |  |
| ATP2B1 | Plasma membrane calcium-transporting ATPase 1 | (Shurtleff et al., 2018) |
| ATP1A1/<br>ATP1B1 | Sodium/potassium-transporting ATPase subunit alpha-1 / beta-1 | (Satoh et al., 2015) |

<sup>a</sup> This table lists previously identified EMC substrates (Satoh et al., 2015; Shurtleff et al., 2018; Tian et al., 2019) that are known regulators of cellular ion homeostasis and cell volume regulation (Jentsch, 2016).

## Acknowledgments

We thank Christian Schlieker for reagents and John O’Neill, Ramanujan Hegde, Henry Lester, and the Voorhees lab for thoughtful discussions. Confocal imaging was performed in the Caltech Biological Imaging Facility with the support of the Caltech Beckman Institute and the Arnold and Mabel Beckman Foundation. This work was supported by the Heritage Medical Research Institute, the Kinship Foundation, the Pew-Stewart Foundation, and the National Institute Of General Medical Sciences of the National Institutes of Health under Award Number DP2GM137412. T.P. is supported by a postdoctoral fellowship from the Deutsche Forschungsgemeinschaft and received prior funding from a Caltech Ross fellowship.

## Author contributions

T.P, A.G, and R.V. conceived the study. T.P., R.O, and M.H. performed the experiments. T.P., R.V., K.J., G.P.T., M.S., and A.M. analyzed the data. T.P. and R.V. wrote the manuscript with input from all authors.

## Material & Methods

### Plasmids, antibodies and siRNAs

Constructs for expression in rabbit reticulocyte lysate (RRL) were based on the SP64 vector (Promega, USA). Constructs for translation in the PURE system were generated from the T7 PURExpress plasmid (New England Biolabs, USA). The coding sequence of full length human WNK1 was derived from a commercial expression vector (#RC218208, Origene, USA). Expression constructs needed to generate biotinylated anti-GFP nanobody for native purification from human cells are available from Addgene (#149336, #149334, #149333) (Pleiner et al., 2015; Pleiner et al., 2020; Vera Rodriguez et al., 2019). Expression constructs for the *bd*NEDP1, *bd*SENP1 and SENP^EuB^ proteases were kind gifts of Dirk Görlich and are available from Addgene (#104129, #104962, #149333) (Frey and Görlich, 2014; Pleiner et al., 2018; Vera Rodriguez et al., 2019). The human NKCC1 coding sequence was derived from Addgene plasmid # 49077, which was a gift from Biff Forbush. The TagBFP coding sequence was derived from Addgene plasmid #67168, which was a gift from Veit Hornung.

The following antibodies were used in this study: WNK1 (A301-515, Bethyl Laboratories, USA); EMC2 (25443-1-AP, Proteintech, USA); EMC3 (67205-1-Ig, Proteintech, USA); EMC4 (27708-1-AP, Proteintech, USA); EMC5 (A305-833, Bethyl Laboratories, USA); EMC6 (ab84902, Abcam, UK); EMC8 (STJ117038, St. John’s Laboratory, UK); Hsp70 (ADI-SPA-812, Enzo Life Sciences, USA); BiP (610978, BD Bioscience, USA); AE2 (A304-502, Bethyl Laboratories, USA); NKCC1 (8351S, Cell Signaling Technologies, USA); CLINT1 (A301-926, Bethyl Laboratories, USA); γ-AP1 (A304-771A, Bethyl Laboratories, USA); FLAG M2 (F1804, Millipore Sigma, USA); Clathrin Heavy Chain (A304-743A, Bethyl Laboratories, USA); β-Actin (8457S, Cell Signaling Technologies, USA); SPAK/OSR1 (S637B, MRC PPU Reagents and Services, United Kingdom); SPAK/OSR1 pSer373/325 (#07-2273, Millipore-Sigma, USA); Anti-HA-HRP (Millipore-Sigma, USA); Anti-FLAG-HRP (Millipore-Sigma, USA). The rabbit polyclonal antibody against BAG6 was previously described (Mariappan et al., 2010). The rabbit polyclonal antibody against GFP was a gift from Ramanujan Hegde. Secondary antibodies used for Western blotting were: Donkey anti-mouse- and anti-rabbit-HRP (#172-1011 and #170-6515, Bio-Rad, USA). Secondary antibodies used for immunofluorescence were: Donkey anti-mouse-Alexa 488 (Invitrogen, USA) and Donkey anti-rabbit-Alexa 647 (Jackson Immunoresearch, USA).

The following siRNAs were used in this study: WNK1 – siRNA (1): On-Targetplus siRNA J-005362-05-0002; siRNA (2): On-Targetplus siRNA J-005362-06-0002 (both from Dharmacon, USA); siRNA (3): s35233; siRNA (4): s35234 and siRNA (5): s35235 (Silencer Select, ThermoFisher Scientific, USA); EMC2 s18670 and EMC5 s41131 (Silencer Select, ThermoFisher Scientific, USA).

### Expression and purification of proteins

To map the EMC2 binding site on WNK1, full-length human WNK1 (residues 1-2382) and its truncations (residues 1-2106, 1-2200, 1-2272) were expressed with a C-terminal Myc-DDK (FLAG) tag from a pCMV6-Entry backbone (#RC218208, Origene, USA) in Expi 293F suspension cells (Thermo Fisher Scientific, USA). Cells were grown in Expi 293 Expression Medium (Thermo Fisher Scientific, USA) at 37°C, 8% CO_2_ and 125 rpm shaking in 1 L roller bottles with vented caps (Celltreat, USA). For a small-scale prep, a 300 ml culture at 2×10^6^ cells/ml was transfected with 330 μg DNA, which was pre-incubated with 900 μl 1 mg/ml PEI MAX 40k (Polysciences, USA) for 15 min at room temperature in 20 ml Expi 293 Expression Medium. Transfected cells were grown for 48-60 hours before harvest. The cell pellet was resuspended and incubated for 30 min in lysis buffer (50 mM HEPES/KOH pH 7.5, 125 mM KAc, 5 mM MgAc_2_, 0.5% (v/v) Triton-X-100, 1 mM DTT and 1x complete EDTA-free protease inhibitor cocktail [Roche, Germany]). After centrifugation (17,000 rpm, 20 min, 4°C, SS-34 rotor, Sorvall, USA), proteins were purified from the lysate with anti-FLAG M2 affinity gel (Millipore-Sigma, USA) and eluted with 0.2 mg/ml 3xFLAG peptide in elution buffer (50 mM HEPES/KOH pH 7.5, 300 mM KAc, 5 mM MgAc_2_, 1 mM DTT, 10% glycerol). Purified DDK(FLAG)-tagged full length WNK1 or C-terminal truncation fragments were added to *in vitro* translation reactions of 3xHA-tagged human EMC2 in RRL. After HA immunoprecipitation and elution with 3xHA peptide, the eluates were analyzed by SDS-PAGE and Western blotting with an anti-FLAG antibody to detect co-purification of WNK1 fragments with EMC2.

Human EMC2, EMC8, EMC3 coiled coil (3-CC; residues 35-117), EMC3 C-terminus (3-CT; residues 196-261), EMC5 C-terminus (5-CT; residues 66-end), and all human WNK helix variants were purified in a similar manner. Open reading frames were fused to N-terminal His_14_-*bd*SUMO or His_14_-*bd*NEDD8 tags (Frey and Görlich, 2014) and expressed in *E. coli* Rosetta 2 (Novagen, USA). 3-CC, 3-CT, 5-CT and all WNK1 helices were additionally fused to a C-terminal 3xFLAG-tag. Typical expressions were carried out in a 1 L scale for 6 hours at 18°C using 0.2 mM IPTG for induction. Cells were resuspended in lysis buffer (50 mM Tris/HCl pH 7.5, 300 mM NaCl, 20 mM imidazole, 1 mM DTT, 1 mM PMSF). Proteins were purified from lysate by Ni^2+^-chelate affinity chromatography and eluted either with imidazole elution buffer (50 mM Tris/HCl pH 7.5, 300 mM NaCl, 500 mM imidazole, 250 mM sucrose) to keep the tag or protease elution buffer (50 nM *bd*SENP1 or 300 nM *bd*NEDP1 in 50 mM Tris/HCl pH 7.5, 300 mM NaCl, 20 mM imidazole, 250 mM sucrose) to remove the N-terminal tag. *bd*NEDP1 and *bd*SENP1 proteases were expressed as described before (Frey and Görlich, 2014; Pleiner et al., 2018). For tagged proteins, the imidazole was removed using a PD-10 desalting column (GE Healthcare, USA).

The expression and purification of GST-3xFLAG-Calmodulin and BpA-RS was carried out as previously described (Shao et al., 2017).

### Expression and purification of biotinylated anti-GFP nanobody

The expression of His_14_-Avi-SUMO^Eu1^-anti GFP nanobody from plasmid pTP396 (Addgene # 149336) was carried out exactly as described in detail before (Pleiner et al., 2020). The anti-GFP nanobody GBP1/Enhancer was described before (Kirchhofer et al., 2010). The Avi-tag is a 15 amino acid peptide with a single lysine that can be specifically biotinylated using purified *E. coli* biotin ligase BirA (Beckett et al., 1999; Fairhead and Howarth, 2015). The expression of Biotin ligase BirA from pTP264 (Addgene #149334) was carried out as described before (Pleiner et al., 2020). The engineered SUMO^Eu1^ module is resistant to cleavage by endogenous eukaryotic deSUMOylases and thus allows stable isolation of nanobody-target complexes from eukaryotic cell lysates, followed by proteolytic release on ice in physiological buffer using the cognate SENP^EuB^ protease (Vera Rodriguez et al., 2019). The expression of SENP^EuB^ protease was carried out as described before (Pleiner et al., 2020; Vera Rodriguez et al., 2019).

### Mammalian *in vitro* translation

*In vitro* translations were carried out in RRL as previously described (Sharma et al., 2010). Briefly, templates for *in vitro* transcription were generated by PCR using primers that anneal 5’ of the SP6 promotor and 3’ of the stop codon of an open-reading frame encoded on either a gene block (Integrated DNA Technologies, USA) or SP64-derived plasmid. Transcription reactions were carried out using SP6 polymerase (New England Biolabs, USA) for 1.5 hours at 37 °C and then added directly to nucleased RRL in the presence of radioactive ^35^S-methionine for 20 min at 32°C. Nascent proteins were subsequently analyzed by SDS-PAGE and autoradiography. For native immunoprecipitation of FLAG-tagged proteins from RRL for analysis by mass spectrometry or Western blot, translations were carried out for 20 min at 32°C. The reactions were then spun for 20 min at 120,000 rpm at 4°C (TLA120.1 rotor, Beckman Coulter, USA) to remove aggregates and ribosomes. Anti-FLAG M2 affinity gel was added to the supernatant and incubated for 1.5 hours at 4°C. After washing with physiologic salt buffer (PSB) (50 mM HEPES/KOH pH 7.5, 130 mM KAc, 2 mM MgAc), bound proteins were eluted with 0.2 mg/ml 3xFLAG peptide in PSB. For analysis of ubiquitination in RRL, translations were performed in the presence of 10 μM HA- or His-tagged Ubiquitin (Boston Biochem, USA). Denaturing immunoprecipitation of the ubiquitinated species was performed as previously described (Yanagitani et al., 2017) using HA- or Ni^2+^-agarose resin, followed by elution with SDS-PAGE sample buffer for analysis by SDS-PAGE and autoradiography.

In order to test assembly of EMC2 with membrane-bound EMC subunits, canine rough ER microsomes (cRMs) were prepared as previously described (Walter and Blobel, 1983) and added to translations in RRL. After incubation for 45 min at 32°C, membranes were pelleted through a 20% (w/v) sucrose cushion, prepared in PSB, for 20 min at 55,000 rpm (TLA-55 rotor, Beckman Coulter, USA). Membrane pellets were carefully resuspended in PSB, mixed with an equal volume of solubilization buffer (2% [w/v] digitonin in PSB) and incubated for 10 min on ice. After a 10 min spin at 55,000 rpm to remove non-solubilized material, the detergent lysate was subjected to FLAG-IP as described above using 0.1% (w/v) digitonin in wash and elution buffers.

### E3 ubiquitin ligase co-purification assay

Detection of co-purifying E3 ubiquitin ligases with EMC2 (Figure 4E) was carried out as follows. ^35^S-methionine-labeled EMC2 was translated in RRL either alone or in the presence of a 3xFLAG-tagged purified binding partner that was not radioactively labeled. Native immunoprecipitations were carried out as described above, utilizing the tag on the binding partner to selectively purify assembled complexes. The immobilized complexes were washed with PSB, and the resin was incubated for 45 min at 32°C with a purified ubiquitination mix that lacked an E3 ligase (PSB containing 10 μM His-tagged Ubiquitin, 1 mM ATP, 40 μg/ml Creatine kinase, 10 mM Creatine phosphate, 100 nM GST-tagged Ubiquitin-activating enzyme E1 and 250 nM Ubiquitin-conjugating enzyme E2 UbcH5 [both from Boston Biochem, USA]). The beads were transferred to 95°C for 5 min in SDS-Buffer (1% [w/v] SDS, 0.1 M Tris/HCl pH 8.4) to terminate ubiquitination and to elute FLAG resin-bound proteins. The samples were then diluted 10-fold with IP-Buffer (50 mM HEPES/KOH pH 7.5, 300 mM NaCl, 0.5% [v/v] Triton-X-100) and subjected to Ni^2+^-chelate affinity chromatography to purify His-ubiquitin-fused proteins. Nickel-bound proteins were eluted with SDS-PAGE sample buffer containing 50 mM EDTA.

Samples of the denaturing FLAG elution that served as input for the Ni^2+^-IP were analyzed first by SDS-PAGE and autoradiography to normalize the amount of purified *in vitro* translated substrate. The resulting normalization was then applied to the Ni^2+^-eluates to load an adjusted SDS-PAGE gel for quantification of ubiquitination via co-purifying E3 ligases.

Alternatively, to analyze co-fractionation of E3 ligases with EMC2 (as in Figure S1C), a 200 μl translation of 3xFLAG-tagged EMC2 in RRL as described above was layered on top of a 2 ml 10-50% (w/v) sucrose gradient in PSB, and centrifuged for 2 hours at 55,000 rpm (TLS-55 rotor, Beckman Coulter, USA). 11x 200 μl fractions were harvested and subjected to FLAG-IP, on-bead incubation with ubiquitination mix as described above and Ni^2+^-chelate affinity chromatography to purify His-ubiquitin-fused proteins.

### *In vitro* translation in the PURE System and photocrosslinking

Site-specific UV crosslinking was carried out as previously described (Shao et al., 2017). A T7 promotor-based construct, encoding human SEC61β with an amber stop codon at position 85 in its TMD, was translated in the presence of radioactive ^35^S-methionine, 100 nM Ca^2+^ and 10 μM purified Calmodulin (CaM) in the coupled transcription/translation PURE system (New England Biolabs, USA), which contains a defined set of purified translation factors and *E. coli* ribosomes. It lacks all release factors (ΔRF). The release factors RF2 and RF3, but not RF1 (which recognizes the UGA [amber] stop codon) were added back to the reaction. The unnatural amino acid 4-Benzoylphenylalanine (BpA, used at 100 μM) was incorporated at the amber stop codon using recombinant BpA synthetase (100 μg/ml) and suppressor tRNA purified as described before (Shao et al., 2017). After translation for 2 hours at 32°C and addition of 1 mM puromycin, the reaction was layered on top of a 20% (w/v) sucrose cushion prepared in PSB with 100 nM CaCl_2_ and spun for 1 hour at 55,000 rpm (TLS-55 rotor, Beckman Coulter, USA) to remove aggregates. SEC61β-CaM complexes were removed from the cushion and incubated in the presence of 1 mM EGTA with either buffer, 3 μM purified EMC2 alone or pre-formed complexes of 3 μM EMC2 and 3 μM of any binding partner (Figure 6B). For Figure 4B, 3 μM EMC2 was pre-incubated with either 3, 6, or 12 μM wild type or L2250K His_14_-*bd*SUMO-tagged WNK1 helix (residues 2241-2266). Except for the -UV control sample, all reactions were irradiated at a distance of ~7-10 cm with a UVP B-100 series lamp (Analytik Jena, Germany) for 15 min on ice before quenching with SDS-PAGE sample buffer. Samples were analyzed by SDS-PAGE and autoradiography.

For the experiment in Figure 6D, a T7-promoter-based construct was generated in which the 3xFLAG-tagged squalene synthase (SQS/FDFT1) transmembrane domain including flanking residues (378-410) was inserted into the inert cytosolic linker region of Sec61β (Guna et al., 2018). It was then translated in the PURE system for 45 min. at 32°C in the presence or absence of 12 μM Calmodulin or 12 μM EMC2. Translation reactions were then layered on top of a 5-25% (w/v) sucrose gradient and spun for 2 hours at 55,000 g. Gradients were fractionated from the top and fractions analyzed by SDS-PAGE and autoradiography.

### Stable cell line generation with the Flp-In system

Flp-In 293 T-Rex cells were purchased from Thermo Fisher Scientific (USA) (RRID: CVCL_U427). HeLa Flp-In cells were a gift from Christian Schlieker. Both cell lines were grown in DMEM supplemented with 2 mM glutamine, 10% FBS, 15 μg/ml Blasticidine S and 100 μg/ml Zeocine. The open-reading frame (ORF) to be integrated into the genomic FRT site was cloned into the pcDNA5/FRT/TO vector backbone and transfected together with pOG44 Flp-In recombinase in a 9:1 ratio using either Trans-IT 293 transfection reagent (Mirus, USA) or Lipofectamine 3000 (Thermo Fisher Scientific, USA) according to the manufacturer’s instructions. 48 hours following transfection, the medium was exchanged to contain 100 μg/ml Hygromycin B to select for cells with a successfully integrated ORF. The resulting cell lines expressed a protein-of-interest from a single genomic integration site under control of a doxycycline-inducible CMV promoter. This procedure was used to generate cell lines stably expressing control GFP-2A-RFP, EMC4-GFP-2A-RFP, EMC5-GFP-2A-RFP, GFP-EMC8-2A-RFP, GFP-NKCC1-2A-RFP, EMC5-3xFLAG, 3xFLAG-WNK1 (wild type) and 3xFLAG-WNK1 (L2250K). Cell lines expressing GFP-EMC2-2A-RFP, TRAM2-GFP-2A-RFP, OPRK1-GFP-2A-RFP as well as the RFP-tagged TMD and flanking regions of human squalene synthase (SQS/FDFT1) or vesicle-associated membrane protein 2 (VAMP2) were previously described (Guna et al., 2018; Pleiner et al., 2020).

### Flow cytometry analysis of reporter cell lines

In order to measure protein stability in cells, stable cell lines were generated as described above containing a fusion of a protein-of-interest with either GFP or RFP in a GFP-2A-RFP backbone (Itakura et al., 2016). Note, the mCherry sequence is used throughout the manuscript, but is referred to as RFP for simplicity. The viral 2A sequence induces the skipping of a peptide bond by the ribosome (de Felipe et al., 2006), resulting in two separate polypeptide chains being produced from a single transcript in a 1:1 ratio. The GFP and RFP fluorescence intensity can be measured by flow cytometry to derive a GFP:RFP or RFP:GFP ratio. Changes in these ratios after perturbation, e.g. siRNA transfection, reflect differences in post-translation stability of the fusion protein. For the analysis of the effect of full length WNK1 (Figure 2A) or WNK1 helix (Figure 3B and 4A) on the stability of EMC2, the respective reporter cell line was transiently transfected with pCDNA3-derived plasmids encoding TagBFP-fusions of these proteins. As a control, a pcDNA3 plasmid encoding only TagBFP was used. The GFP:RFP ratio of only BFP-positive cells was then compared.

All cells for flow cytometry were trypsinized, washed in 1xPBS, dissolved in Hank’s balanced salt solution containing 50 μg/ml DNAse I and analyzed using a MACSQuant VYB flow cytometer (Miltenyi Biotec, Germany). Sytox Blue dead cell stain (ThermoFisher Scientific, USA) was used at 2 μM, where possible, to gate out dead cells. Unstained wild type Flp-In cells and Flp-In cells transiently transfected with either GFP, RFP (or BFP if needed) were analyzed separately along every run as single-color controls for multicolor compensation using the FlowJo software package.

### Affinity purifications from stable cell lines

To validate incorporation of GFP-tagged subunits into the EMC, we isolated the EMC under native conditions from detergent-solubilized cells using a biotinylated, protease-cleavable anti-GFP nanobody, expressed and purified as described above (Pleiner et al., 2020). Cells were grown in 15 cm dishes and GFP-EMC2/4/5/8 expression was induced for at least 48 hours with 1 μg/ml DOX. Cells were then harvested by centrifugation, washed with 1x PBS, weighed and resuspended with 6.8 μl solubilization buffer per 1 mg cell pellet (50 mM HEPES/KOH pH 7.5, 200 mM NaCl, 2 mM MgAc_2_, 1% [w/v] LMNG, 1 mM DTT, 1x complete EDTA-free protease inhibitor cocktail [Roche, Germany]). After 30 min. of head-over-tail incubation with solubilization buffer, the lysate was cleared by centrifugation for 20 min at 4°C and 18,000 x g in a tabletop centrifuge. The supernatant was then added to pre-equilibrated magnetic Streptavidin beads (ThermoFisher Scientific, USA) onto which biotinylated anti-GFP nanobody had been pre-immobilized before blocking all free binding sites with biotin. After 1 hour of binding head-over-tail at 4°C, the beads were washed three times with wash buffer (solubilization buffer with 0.1% [w/v] LMNG). Finally, the anti-GFP nanobody along with all bound proteins was released under native conditions and in minimal volume by cleavage with 250 nM SENP^EuB^ protease (Vera Rodriguez et al., 2019) in wash buffer for 20 min at 4°C. The eluate was then analyzed by SDS-PAGE and Western blotting or Sypro Ruby staining (Bio-Rad Laboratories, USA).

Alternatively, the EMC was purified from HEK293T cells stably expressing human EMC5-3xFLAG as previously described (Guna et al., 2018).

### Analysis of EMC subunit mutations in cells

To test incorporation of mutant EMC subunits into intact EMCs in cells (Figures 5E-F and S6I), lenti-viral vectors in a pHAGE2 backbone were generated that stably integrate and express either wild type or mutant subunit from a CMV promotor-driven open-reading frame. EMC2 variants were fused to an N-terminal 3xFLAG-TagBFP tag, while EMC5 variants carried a C-terminal TagBFP-3xFLAG tag. A 2^nd^ generation packaging system was used to generate lenti-viral particles. Cells were transduced with lenti-viral particles such that around 20 % of all cells were BFP-positive, likely indicating integration of a single copy of the transgene. After 48 hours, cells were harvested, solubilized as described above and subjected to native FLAG-IP. Bound proteins were eluted with 3xFLAG peptide and the eluates analyzed by SDS-PAGE and Western blotting.

### Immunofluorescence and confocal microscopy

Lab-Tek II 8-well chamber slides (ThermoFisher Scientific, USA) were coated with 300 μl 100 μg/ml poly-L-lysine per well for 1.5 hours at 37°C and then washed three times with ddH_2_O. Stable HEK293T cells expressing 3xFLAG-tagged wild type or L2250K mutant WNK1 were seeded onto coated slides 24 hours before analysis in medium containing 1 μg/ml DOX. For the hyper-osmotic shock, the medium was aspirated and exchanged to new medium additionally containing 200 mM sorbitol for 5 min. Then all medium was aspirated and cells fixed with 4% (w/v) paraformaldehyde in 1x PBS for 15 min at room temperature. After two washes with 1x PBS, a 5 min quenching step with 50 mM NH_4_Cl in 1x PBS and two further washes with 1x PBS, cells were permeabilized with 0.2% (v/v) Triton-X-100 in 1x PBS for 3 min. Following three washes with 1x PBS, cells were blocked for 30 min with 1% (w/v) bovine serum albumin fraction V (BSA) in 1x PBS. Then cells were incubated with anti-FLAG M2 antibody (1:500) and anti-Clathrin heavy chain antibody (1:100) diluted in 1x PBS containing 1% (w/v) BSA. After three 10 min washes with 1x PBS, cells were stained with the respective fluorophore-coupled secondary antibodies and 2.5 μg/ml DAPI diluted in 1x PBS containing 1% (w/v) BSA. Following another three 10 min washes with 1x PBS, cells were mounted for analysis by confocal microscopy using SlowFade Gold (Thermo Fisher Scientific, USA). Imaging was performed using a LSM880 confocal microscope (Zeiss, Germany).

---

**Figure S1.**
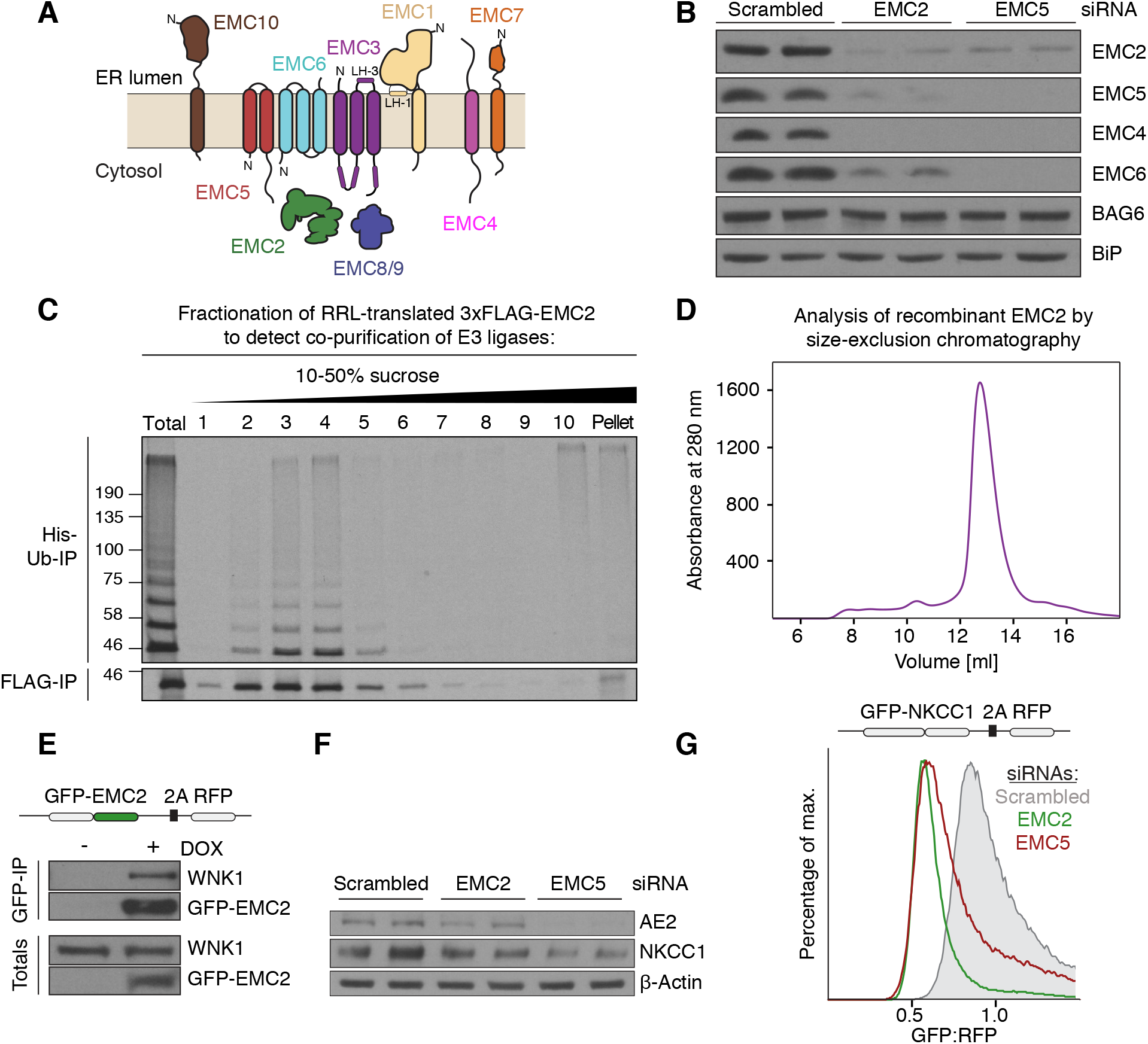
Unassembled EMC2 is soluble, monodisperse and recognized by the ubiquitin proteasome pathway. (**A**) Topology of the EMC subunits based on the structure of the human EMC (Pleiner et al., 2020). Note that EMC8 and 9 are paralogs and their binding to EMC2, and thus incorporation into the EMC, is mutually exclusive. (**B**) The stability of soluble and membrane-embedded EMC subunits is interdependent in cells. HeLa cells were treated with scrambled, EMC2, or EMC5 siRNA and harvested for Western blotting with the indicated antibodies. Similar results were observed in HEK293T, U2OS, and RPE1 cell lines and has been reported in previous studies (Guna et al., 2018; Volkmar et al., 2019). (**C**) Soluble EMC2 stably recruits an E3 ubiquitin ligase. An *in vitro* translation reaction of ^35^S-methionine-labeled 3xFLAG-EMC2 was fractionated on a 10-50% (w/v) sucrose gradient. Unfractionated translation (Total) and all fractions were subjected to FLAG-IP and subsequently incubated with ATP, His-tagged ubiquitin (His-Ub), as well as E1 and E2 enzymes to detect the presence of co-purifying E3 ligase activity. Ubiquitinated species were enriched after elution from the FLAG resin by Ni^2+^-chelate affinity chromatography. (**D**) *E. coli*-purified EMC2 is a monodisperse, soluble protein. His_14_-*bd*NEDD8-tagged EMC2 was purified on a Superdex 200 Increase 10/300 size-exclusion chromatography column. UV absorbance at 280 nm was monitored throughout the purification. (**E**) WNK1 co-purifies with EMC2 from human cells. HEK293T cell lines stably expressing GFP-EMC2 were used for purification of GFP-tagged EMC2 and its interacting proteins under native conditions using a biotinylated Avi-SUMO^Eu1^-tagged anti-GFP nanobody (see methods). Samples of total lysate and eluate were analyzed by Western blot with anti-GFP and anti-WNK1 antibodies. (**F**) Endogenous NKCC1 expression relies on a functional EMC. HeLa cells were treated with either scrambled, EMC2, or EMC5 siRNA and harvested for Western blotting with the indicated antibodies. Expression of endogenous AE2, a previously identified EMC substrate, displays a similar dependence on the EMC for expression. (**G**) NKCC1 is post-translationally destabilized in the absence of the EMC. HEK293T cells stably expressing GFP-NKCC1-2A-RFP were treated with either scrambled, EMC2, or EMC5 siRNA and the GFP:RFP ratio was analyzed by flow cytometry. The observed post-translational destabilization of NKCC1 upon EMC depletion is consistent with a requirement for EMC in NKCC1 biogenesis.

**Figure S2.**
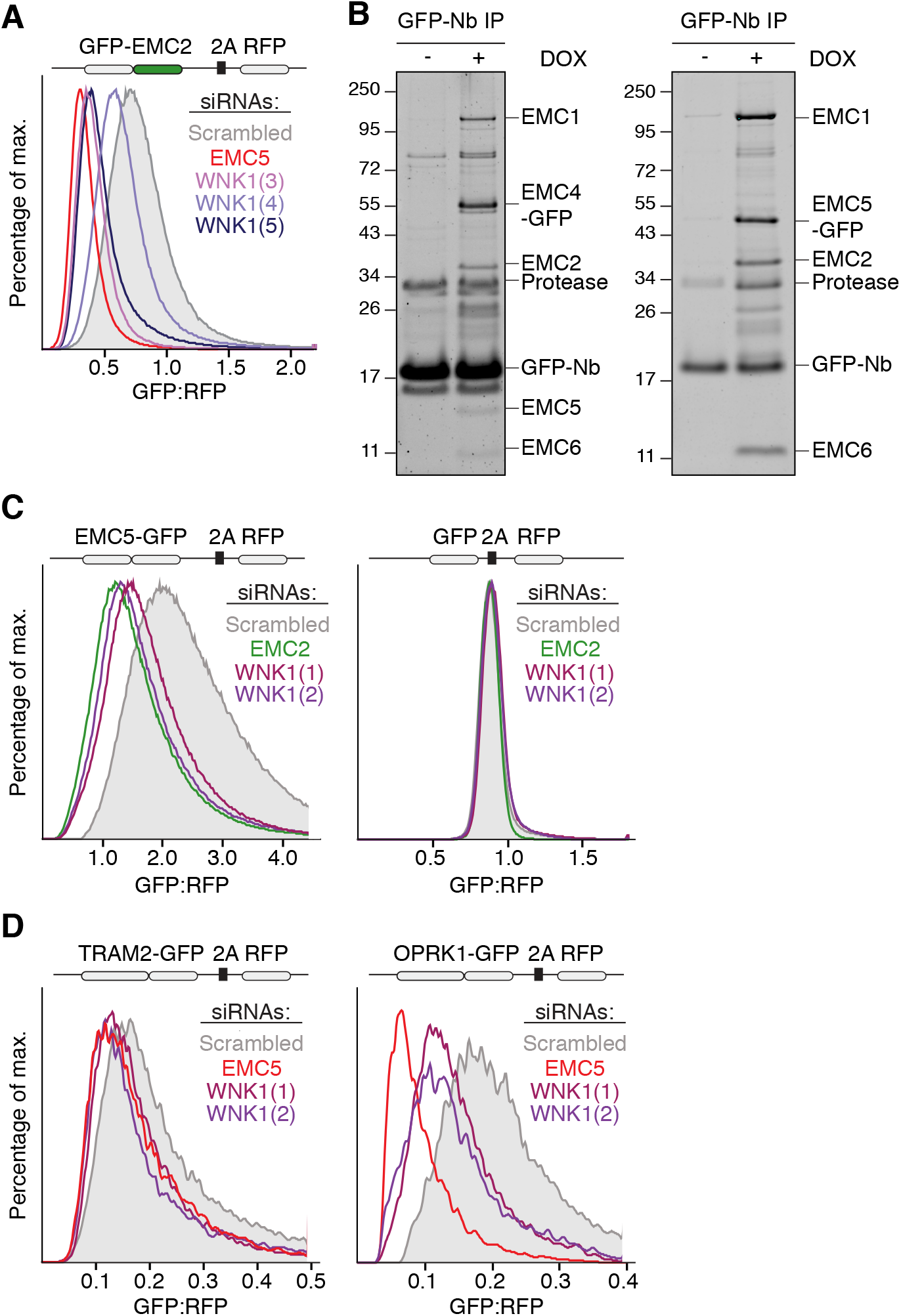
WNK1 knockdown post-translationally destabilizes the EMC and its substrates, consistent with a role in EMC assembly. (**A**) HEK293T cells stably expressing GFP-EMC2-2A-RFP were treated with a scrambled control or EMC5 siRNA, or three independent WNK1 siRNAs for 72 hours. 24 hours following siRNA transfection the fluorescent reporter was induced for 48 hours. Cells were then analyzed by flow cytometry and their GFP:RFP ratio plotted as a histogram. (**B**) HEK293T cell lines stably expressing (left) EMC4-GFP-2A-RFP, and (right) EMC5-GFP-2A-RFP, were harvested 48 hours after reporter induction with doxycycline (DOX), solubilized in 1% (w/v) LMNG, and the GFP-tagged subunits purified using an anti-GFP nanobody (Nb) (see methods) (Pleiner et al., 2020). The eluates were analyzed by SDS-PAGE and Sypro Ruby staining. GFP-tagged EMC4 and 5 are incorporated into the intact EMC and thus co-purify endogenous EMC subunits, which were identified by mass spectrometry. The characterization of the stable HEK293T GFP-EMC2-2A-RFP cell line was described before (Pleiner et al., 2020). (**C+D**) Stable HEK293T cell lines expressing either (**C**, left) EMC5-GFP-2A-RFP or (**C**, right) GFP-2A-RFP without any fusion partner, (**D**, left) TRAM2-GFP-2A-RFP or (**D**, right) OPRK1-GFP-2A-RFP were treated with the indicated siRNAs for 72 hours. 24 hours following siRNA transfection the fluorescent reporter was induced for 48 hours with DOX. Cells were then analyzed by flow cytometry and their GFP:RFP ratio plotted.

**Figure S3.**
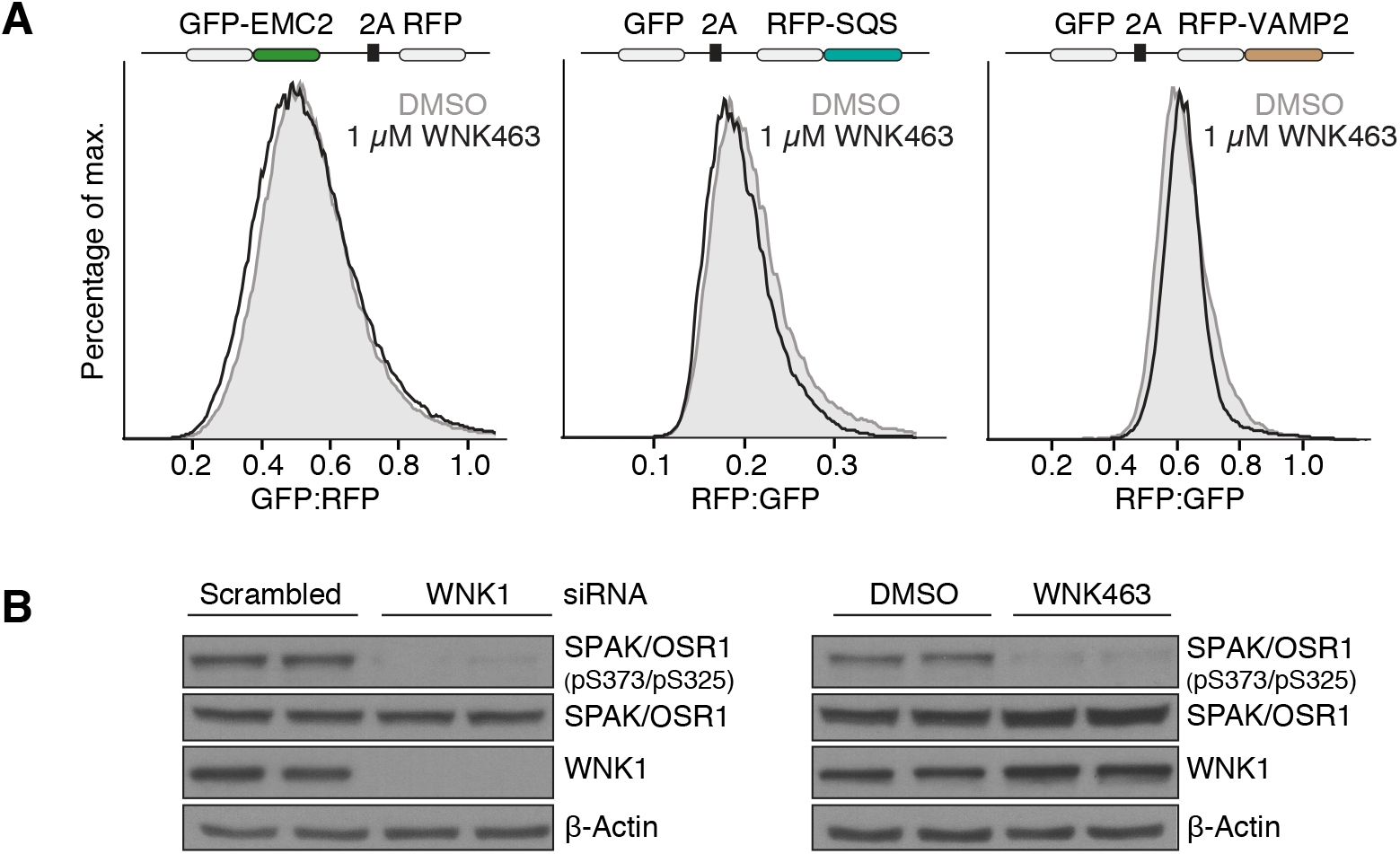
The kinase activity of WNK1 is not required for its role in EMC assembly. (**A**) Stable HEK293T cell lines expressing either GFP-EMC2-2A-RFP (left), GFP-2A-RFP-SQS (middle) or GFP-2A-RFP-VAMP2 (right) were treated either with DMSO or 1 *μ*M pan-WNK inhibitor WNK463 (Yamada et al., 2016) (IC_50_= 5 nM for WNK1) for 68 hours. 20 hours following drug treatment the fluorescent reporter was induced for 48 hours with DOX. Cells were then analyzed by flow cytometry and their GFP:RFP ratio plotted as a histogram. No effect of WNK463 on EMC2 stability or EMC activity was observed. (**B**) Left: HEK293T cells were treated with either scrambled or WNK1 siRNA for 72 hours. Right: HEK293T cells were treated with DMSO or 1 *μ*M WNK463 for 68 hours. Cells were then harvested for Western blotting with the indicated antibodies. Note that WNK463 treatment reduced SPAK/OSR1 S-motif phosphorylation (pSer373 in SPAK, pSer325 in OSR1) much like a WNK1 knockdown.

**Figure S4.**
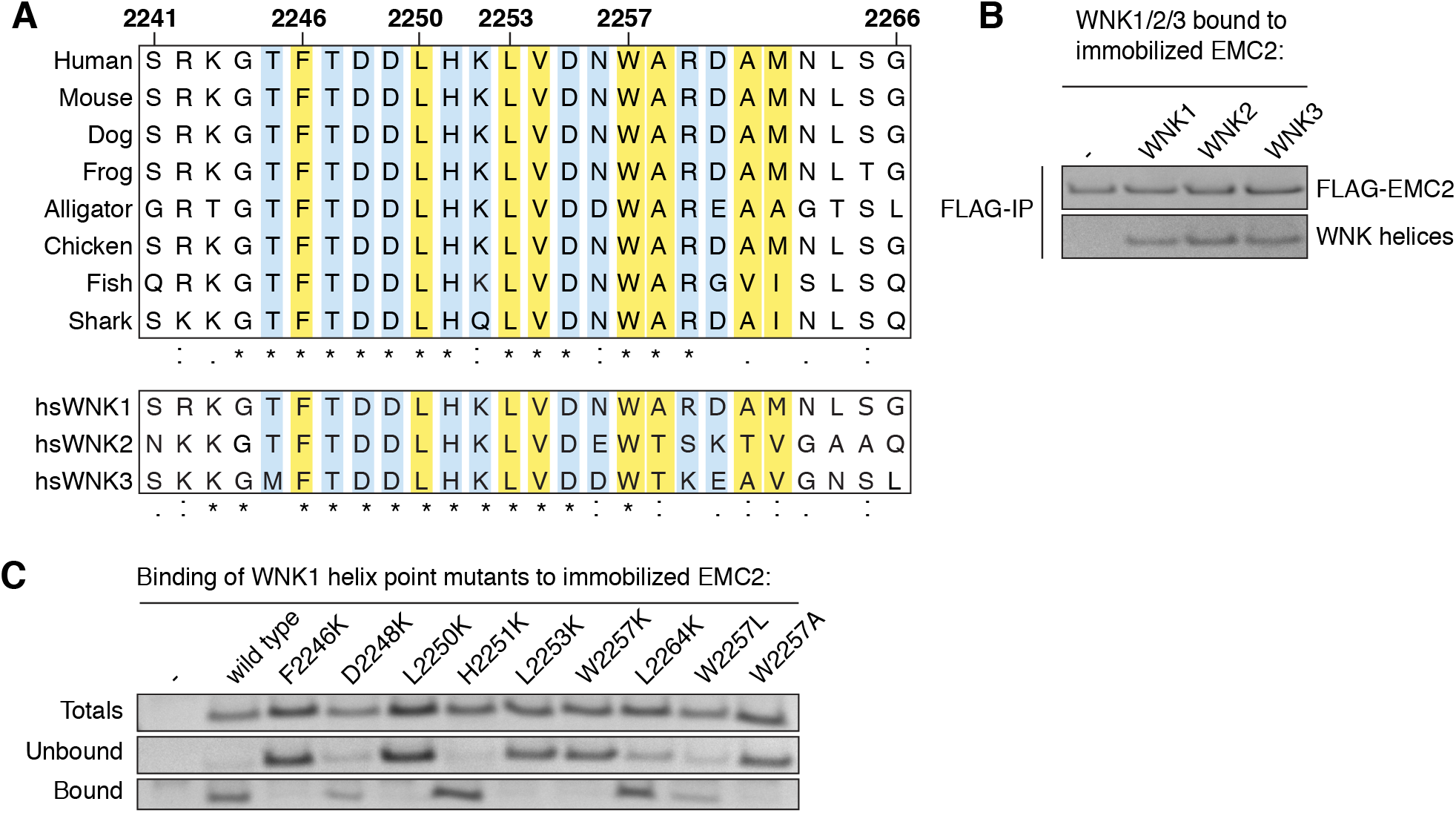
WNK1 amphipathic helix is conserved and binds EMC2 via its hydrophobic face. (**A**) High sequence conservation of the WNK1 helix in vertebrates. Top: Shown are sequence alignments corresponding to residues 2241-2266 in human WNK1. The following species were included: Human (*Homo sapiens*), Mouse (*Mus musculus*), Dog (*Canis lupus familiaris*), Frog (*Xenopus tropicalis*), Alligator (*Alligator sinensis*), Chicken (*Gallus gallus*), Fish (*Nothobranchius furzeri*) and Shark (*Callorhinchus milii*). Bottom: Alignment of human WNK1, WNK2 and WNK3 in the region corresponding to the amphipathic α-helix (residues 2241-2266 in human WNK1). WNK4 has no region with significant sequence homology. (**B**) The equivalent helical regions in WNK2 (residues 2153-2178) and WNK3 (residues 1641-1666) also interact with EMC2. Purified 3xFLAG-tagged EMC2 was incubated with either buffer or His_14_-*bd*SUMO-tagged WNK helix variants and then subjected to FLAG-IP. Bound proteins were eluted with SDS sample buffer and analyzed by SDS-PAGE and Coomassie staining. (**C**) The hydrophobic face of the WNK1 amphipathic helix interacts with EMC2. Purified 3xFLAG-tagged EMC2 was incubated with either buffer or His_14_-*bd*SUMO-tagged WNK1 helix variants and then subjected to FLAG-IP. Bound proteins were eluted with SDS sample buffer. Samples of totals, unbound and elution were analyzed by SDS-PAGE and Coomassie staining.

**Figure S5.**
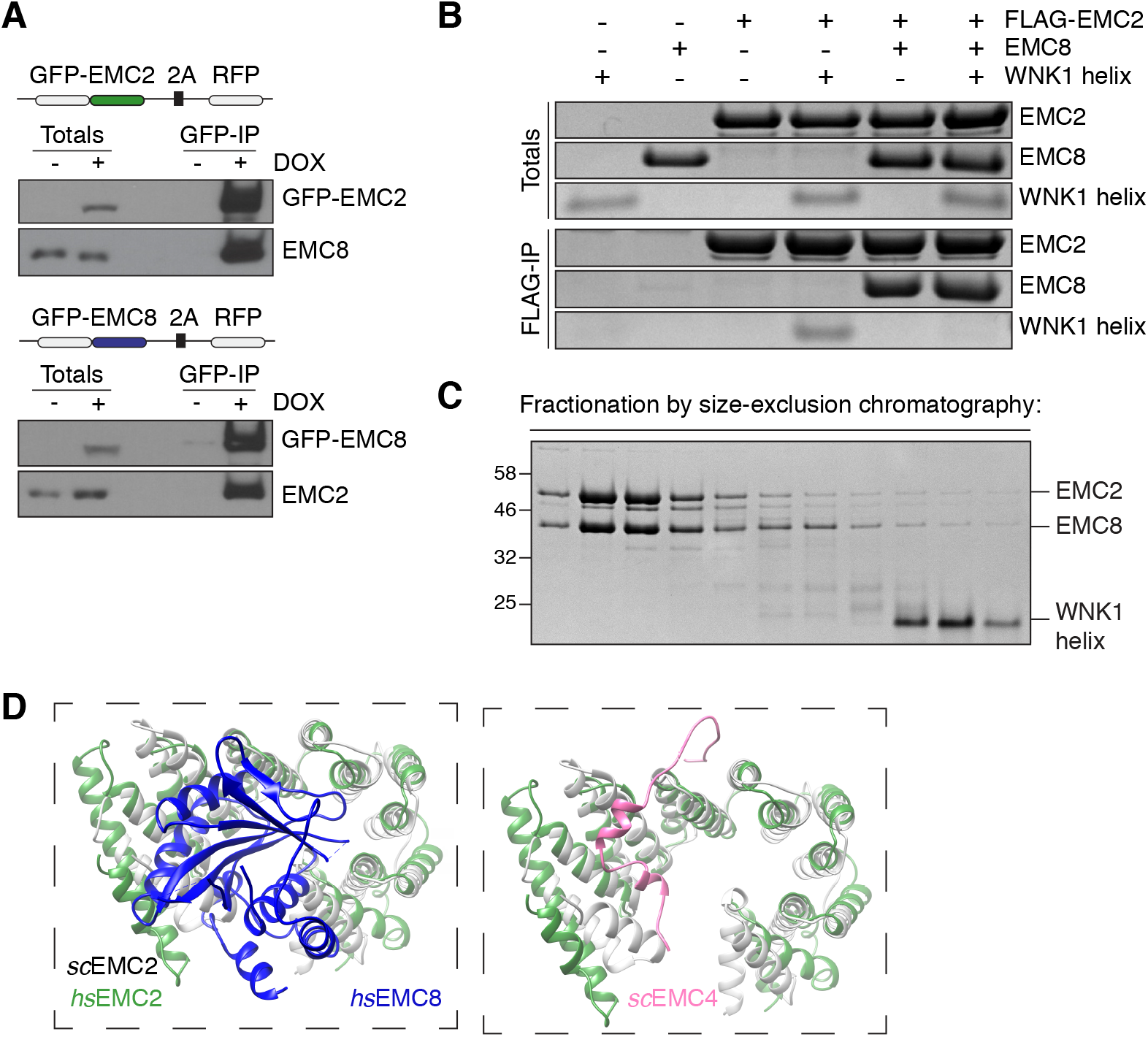
WNK1 binding is mutually exclusive to EMC2-8 complex formation. (**A**) EMC2 and EMC8 form a complex in human cells. Stable HEK293T cell lines were induced with DOX to express GFP-tagged EMC2 or EMC8. After mechanical lysis under non-solubilizing conditions, GFP-tagged EMC2 or EMC8 and their interacting proteins were purified under native conditions from lysate using a biotinylated anti-GFP nanobody (see methods). Samples of total lysate and eluate were blotted and probed with the indicated antibodies. (**B+C**) WNK1 helix binding to EMC2 is incompatible with EMC8 interaction. (**B**) Purified 3xFLAG-tagged EMC2 was incubated either with purified EMC8, WNK1 helix, or both and then subjected to FLAG-IP. Totals and eluates were analyzed by SDS-PAGE and Coomassie staining. WNK1 helix binding to EMC2 is mutually exclusive to binding of EMC8. (**C**) Stoichiometric amounts of EMC2, EMC8 and WNK1 helix were incubated to allow complex formation and then analyzed by size-exclusion chromatography. Selected fractions were analyzed by SDS-PAGE and Coomassie staining. Note that EMC8 outcompetes WNK1 helix binding to EMC2 (compare with Figure S6A). (**D**) In yeast, the analogous WNK1/EMC8 interface on EMC2 is occupied by the N-terminus of EMC4. A structural overlay of yeast EMC2 (gray: PDB 6WB9) and human EMC2 (green: PDB 6WW7) is shown with or without human EMC8 (blue) and yeast EMC4 N-terminus (pink) (Bai et al., 2020; Pleiner et al., 2020). The N-terminus of EMC4 is poorly conserved, consistent with its non-essential role in binding EMC2 in metazoans.

**Figure S6.**
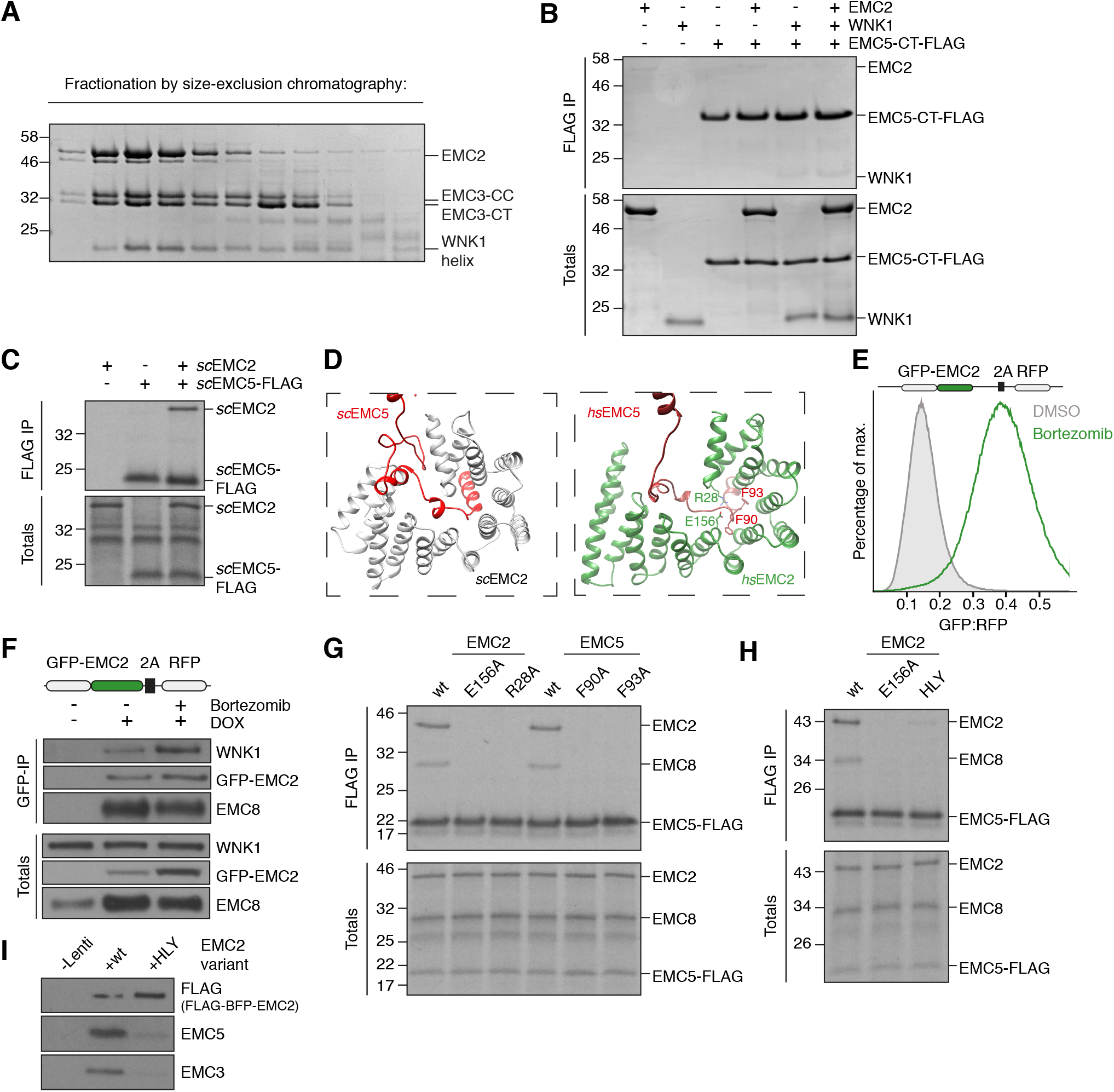
Characterization of the EMC2-3 and EMC2-5 interactions. (**A**) WNK1 helix binding to EMC2 is compatible with EMC3 interaction. Purified EMC2 was incubated with EMC3 coiled coil (3-CC), EMC3 C-terminus (3-CT), and WNK1 helix, and the resulting complexes were analyzed by size-exclusion chromatography. Peak fractions were analyzed by SDS-PAGE and Coomassie staining. (**B**) The EMC2-WNK1 complex is not capable of binding to the EMC5 C-terminus (5-CT). As in Figure 5D, using purified EMC2 and 5-CT, but with WNK1 helix instead of EMC8. (**C**) In the absence of an EMC8 homolog in yeast, *S. cerevisae* (*sc*) EMC2 binds *sc*EMC5 directly. Full-length *sc*EMC5-3xFLAG was translated alone or co-translated with 3xHA-tagged *sc*EMC2 in RRL in the presence of cRMs and subjected to FLAG-IP as described in Figure 5A. Bound proteins were eluted with 3xFLAG peptide and analyzed alongside samples of total translation by SDS-PAGE and autoradiography. (**D**) Comparison of the yeast (PDB 6WB9) and human EMC2-5 interfaces (PDB 6WW7) (Bai et al., 2020; Pleiner et al., 2020). In contrast to human EMC2, the N-terminus of yeast EMC2 packs against an elongated unstructured loop of yeast EMC5, resulting in a more extensive interface. Interface residues in human EMC2 and EMC5 that were mutated in (G) are highlighted. (**E**) EMC2 strongly accumulates upon proteasome inhibitor treatment. HEK293T cells stably expressing GFP-EMC2-2A-RFP were induced with DOX and treated with either DMSO or 20 nM proteasome inhibitor bortezomib. Cells were were analyzed by flow cytometry and their GFP:RFP ratio plotted as a histogram. (**F**) Unassembled EMC2 preferentially engages WNK1 in human cells. Cells were treated as in (E) and then lysed mechanically under non-solubilizing conditions, GFP-tagged EMC2 and its interacting proteins were purified from lysate with biotinylated anti-GFP nanobody. Normalized samples of total lysate and eluate were analyzed by Western blotting with the indicated antibodies. Following bortezomib treatment WNK1 co-purification with EMC2 increases, whereas EMC8 co-purification decreases. This indicates that under conditions where EMC8 becomes limiting, EMC2 accumulates in complex with WNK1. (**G**) ^35^S-methionine-labeled 3xFLAG-tagged EMC5 (or the indicated mutants) was translated in the presence of 3xHA-tagged EMC2 (or the indicated mutants) and EMC8 in RRL in the presence of canine rough microsomes (cRMs). cRMs were isolated by centrifugation through a sucrose cushion, resuspended, and solubilized in digitonin. The resulting detergent lysate was subjected to FLAG-IP and analyzed by SDS-PAGE and autoradiography. (**H**) As in G. EMC2 HLY is a triple mutant of H189E, L190E and Y191K that was previously described (O’Donnell et al., 2020). (**I**) As in Figure 5E, HEK293T cells stably expressing exogenous 3xFLAG-BFP-tagged wild type or the HLY mutant EMC2 were lysed and subjected to native FLAG-IP. Eluates were analyzed by Western blotting.

**Figure S7.**
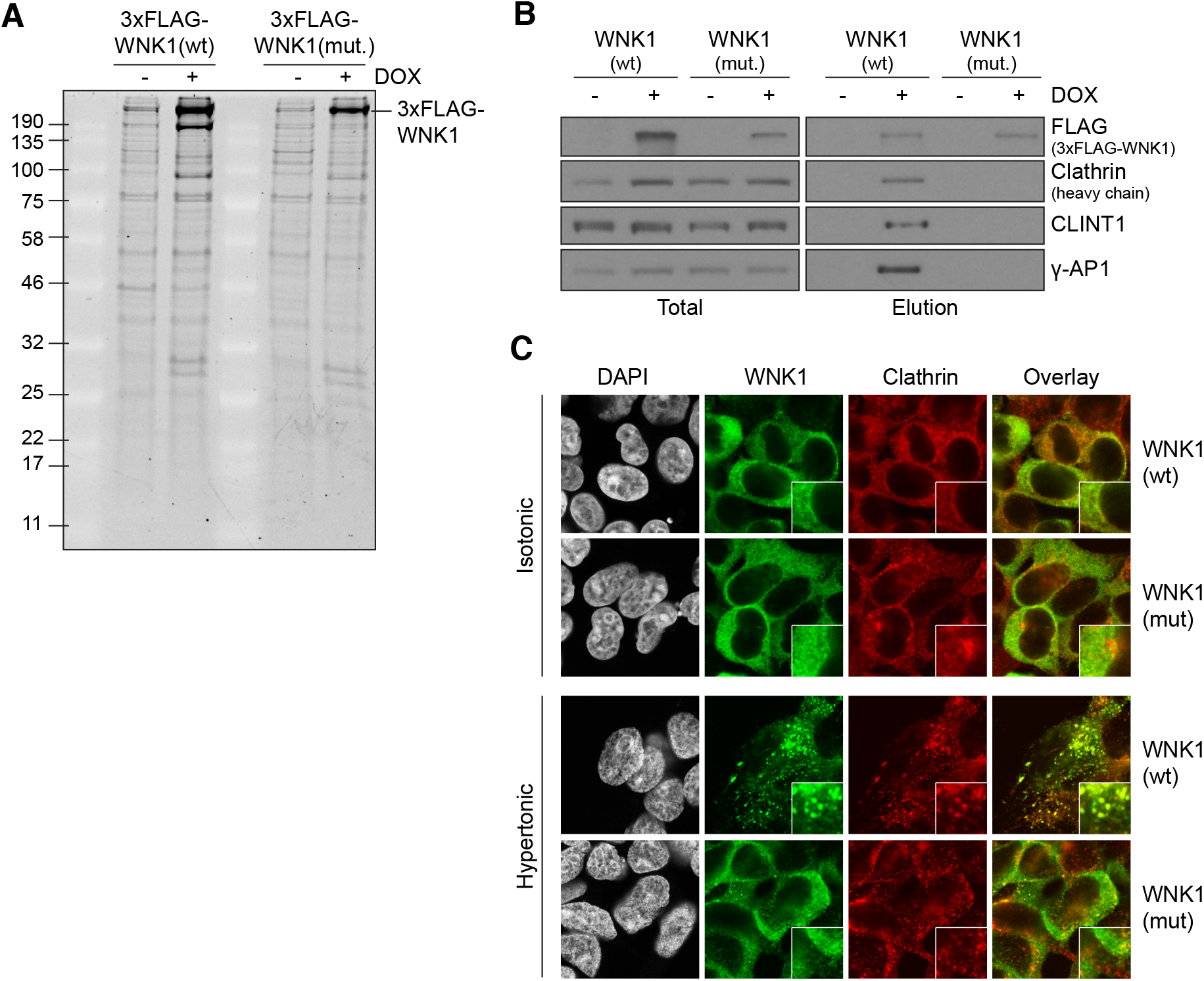
WNK1 amphipathic helix mediates vesicle recruitment. (**A**) WNK1 co-immunoprecipitates with vesicle proteins. Stable HEK293T cell lines expressing 3xFLAG-tagged full length WNK1 wild type (wt) or WNK1 helix mutant (mut) (L2250K, L2253K, W2257K) were lysed mechanically under non-solubilizing conditions. WNK1 and its interacting proteins were immunoprecipitated from lysate with anti-FLAG resin. After elution with 3xFLAG peptide, the eluates were analyzed by SDS-PAGE and Sypro Ruby staining. (**B**) Eluates from (A) were analyzed by Western blotting with the indicated antibodies. Mass spectrometry revealed Clathrin and Clathrin-associated vesicle proteins as the highest specific hits. Importantly, these interactions are sensitive to detergents commonly used for cell lysis (e.g. 1% Triton-X-100 or NP-40), explaining why they were not observed before (Zagórska et al., 2007). Mutation of WNK1’s amphipathic helix disrupts its interaction with vesicle-resident proteins. (**C**) HEK293T cell lines stably expressing full-length 3xFLAG-WNK1(wt) or WNK1 (mut = L2250K, L2253K, W2257K) were either left untreated (isotonic) or stimulated with 0.2 M Sorbitol for 5 min prior to fixation. Cells were then stained with anti-FLAG and Clathrin antibodies via indirect immunofluorescence and analyzed by confocal microscopy.

